# Population genomics of the *Anopheles farauti* complex reveals a distinct regime of malaria vector evolution in the southwest Pacific

**DOI:** 10.64898/2026.11.24.767905

**Authors:** Tristan P. W. Dennis, Luke Ambrose, Fitsum Bekele, Loïc Talignani, Weng K. Chow, Tanya Russell, Michael J. Bangs, Rebecca Vinit, Michelle Katusele, Lincoln Timinao, Solomon Lagur, Petrina Johnson, John Bosco Keven, Andrew F. van den Hurk, Greg Devine, Anthony W. Sweeney, Stephen P. Frances, Darren Watterson, Thomas R. Burkot, Hugo Bugoro, Stephan Karl, Robert D. Cooper, Michael C. Fontaine, Nigel W. Beebe

## Abstract

The southwest Pacific has one of the highest per-capita malaria burdens outside Africa, yet how its malaria mosquitoes evolve is largely unknown. In African vectors, insecticide resistance evolves rapidly in large, connected populations carrying abundant standing variation. Southwest Pacific vectors show little to no biochemical resistance despite decades of insecticide-based control. We analyse 495 genomes from six taxa of the *Anopheles farauti* complex, which includes the region’s major vectors. The complex radiated rapidly in the late Pleistocene concomitant to the peopling of Near Oceania. Thereafter, post-glacial sea-level rise and climate change fragmented its populations into recently colonised island isolates. Successive environmental and colonisation bottlenecks left a legacy of extraordinary inbreeding, most marked in island populations of the coastal specialist *An. farauti* sensu stricto, where sustained low effective population size has allowed deleterious alleles to be purged effectively; the inland generalists *An. hinesorum* and *An. oreios* retain more variation. After five decades of chemical control we recover a single resistance sweep, at *Gste2*, alongside recent selection at species-specific chemosensory gene repertoires. Severe inbreeding and the loss of standing variation have not prevented ecological success, challenging assumptions about how much variation and connectivity a vector population needs to thrive. This work establishes the first genomic foundation for this major vector system of endemic species through a biogeographically diverse region of the world.

## Main Text

Control of malaria, a disease that kills over 500,000 yearly, relies largely on killing vectors using insecticide-treated nets (ITNs) and indoor residual spraying (IRS) of house walls. These tools target the specialised feeding behaviour of anopheline vectors that bite indoors, at night. In response to this anthropogenic selection pressure, *Anopheles* mosquitoes have shown an extraordinary ability to resist chemical control. In Africa - home to the most intensively studied malaria vectors - resistance arises through target-site mutations mostly drawn from standing genetic variation (*1*), and structural variation at metabolic resistance gene families (*2*), leaving strong signatures of positive selection in which favoured alleles spread rapidly across vast spatial scales through large, highly inter-connected populations (*3*, *4*).

The southwest Pacific, where the malaria burden is concentrated in New Guinea and Solomon Islands and per-capita transmission is among the highest outside Africa (*5*), presents a markedly different paradigm under the same interventions (*6*). Malaria is vectored by three dominant species complexes – *An. farauti, An. koliensis,* and *An. punctulatus –* each containing multiple cryptic taxa, alongside numerous secondary vectors (*7–10*). Physiological resistance has rarely been detected (*11*), with only occasional focal reports (*12*). In the absence of physiological resistance, the candidate explanations for sub-optimal control efficacy are unclear. They include operational factors: reduced net efficacy and durability (*13*) and variation in coverage; the behaviour of the vectors themselves, which often bite early in the evening and outdoors (*10*, *14*), and readily feed on non-human hosts (*6*, *15*, *16*). Whether the behavioural component reflects a heritable adaptation to control (*17–19*), turnover in species composition as more endophilic vector species are removed (*20*, *21*), or plasticity already present in these populations, is unclear. Available observations often come from small numbers of villages over short time-scales, and the earliest and most frequently cited examples (*14*) predate molecular assays that can resolve species complexes into their constituent members. Separating them requires knowing how these taxa are related, how their populations are structured, and how they have evolved, and much of that remains unknown.

In 1985 the Australian Defence Force (ADF) embarked on *Operation Anopheles*: a decade-long series of mosquito surveys with helicopter, fixed wing and four wheel drive support to characterise the vectors transmitting malaria in the southwest Pacific. These systematic surveys generated more than 100,000 *Anopheles* occurrence records and an archive of specimens spanning the region (*22–24*). They underpinned malaria vector species identification using molecular species diagnostics (*6*) and early population genetics studies of its complex, highly structured populations (*25–28*) including those of its two most widespread constituent taxa, *Anopheles farauti sensu stricto (= Anopheles farauti)* and *Anopheles hinesorum* (formally *Anopheles farauti* No. 2), both members of the *An. farauti* complex, a group of morphologically indistinguishable species. These data suggested that New Guinea is the centre of diversity from which populations in northern Australia, Solomon Islands and Vanuatu derive (*26*), and pointed to polyphyletic relationships and evidence of introgression events (*26*, *29*) between anthropophilic and zoophilic populations of *An. hinesorum*, and selection at olfactory gene loci (*30*) that may underlie the loss and re-emergence of human-biting in the Western Province of Solomon Islands (*31*). But this earlier work relied on a small number of mitochondrial and nuclear loci and likely overlooked a far more complex evolutionary history. However, despite initial work and substantial interest, this group of vectors remain largely understudied at the genomic level, despite their status as major malaria vectors in the region. This is in stark contrast to the depth of genomic resources now available for their counterparts in Africa (*3*, *4*, *32*), Asia (*33–35*) and the Americas (*36*). In terms of our understanding of vector ecology, adaptation, and evolution, they are the last frontier among the world’s major malaria vectors.

Here, we address this critical gap with a foundational genomic assessment of the *An. farauti* complex, the most geographically widespread and medically important vector group in the region (*6*, *8*) spanning New Guinea, northern Australia, the Solomon Archipelago, and Vanuatu (Fig. 1A). Drawing on archived material from the original ADF surveys, alongside additional and contemporary collections (Fig. 1B, supplementary materials), we sequenced the whole genomes of more than 495 samples from six of the eight species of the *An. farauti* complex. We focus on *An. farauti*, a coastal species whose larvae have a preference for brackish habitats, and *Anopheles hinesorum*, with a more generalist habitat preference, inhabiting lowland mainland regions (*6*) (Fig. 1A), both species are competent vectors and the most widely distributed of all the members of the complex (Fig. 1A). We also included *Anopheles oreios:* a sister species of *An. hinesorum* which only inhabits highland regions (*37*) (Fig. 1A); *Anopheles irenicus:* a non-human biting mosquito native only to the island of Guadalcanal, Solomon Islands; *Anopheles farauti* 8: a minor vector native to the Gulf of New Guinea, and *Anopheles torresiensis*, a species native to the highly monsoonal region of northern Australia and southern New Guinea and whose vector status is unknown (Fig. 1A). *Anopheles punctulatus,* itself a major vector in the region, served as an outgroup for genetic analyses (tables S1 and S2; Fig. 1A).

**Fig. 1.**
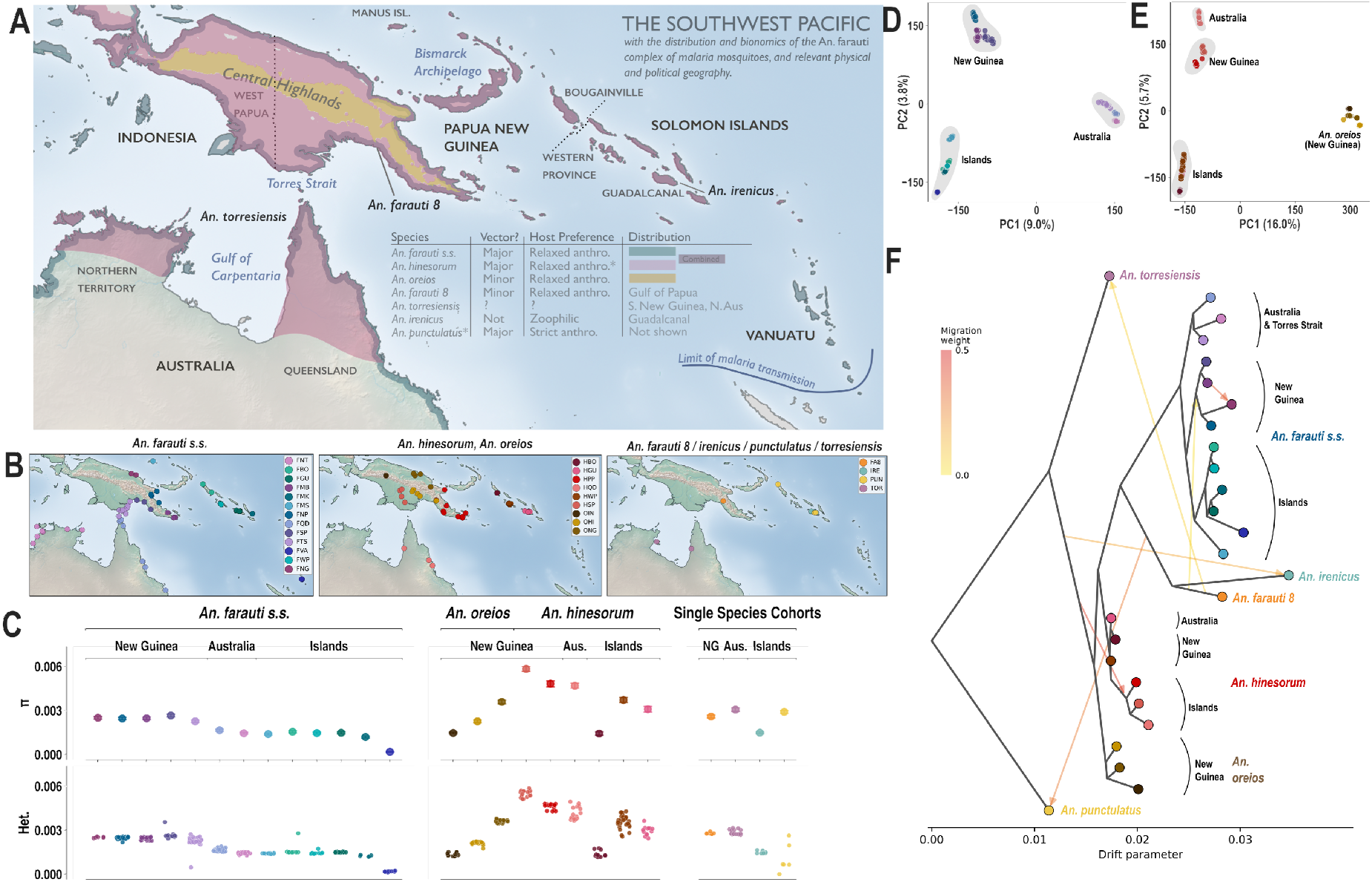
The distribution, population structure and genomic diversity of the *An. farauti* complex. **(A)** Shows the Southwest Pacific, indicating relevant political and geographic features to this study. Also shown are the distribution of the three principal vectors of the *An. farauti* complex: *An. farauti s.s., An. hinesorum* and *An. oreios.* *Note that *An. hinesorum* is mostly strictly zoophilic in the Solomon Archipelago, and relaxed anthropophilic in New Guinea and Australia. *An. irenicus, An. torresiensis* and *An. farauti* 8 are minor or non-vectors highly localised to Guadalcanal, the coasts of Australia and PNG bordering the Carpentaria Gulf and Arafura Sea, and the Gulf of Papua, and are not fully shown but indicated. The full distribution of *An. punctulatus*, outgroup taxon (*6*) is also not shown here. A summary table of vector bionomics is inset. (**B**) shows sampling locations and population definitions (**see** supplementary materials and table S2) of *An. farauti s.s., An. hinesorum* and *An. oreios,* and *An. irenicus, punctulatus, torresiensis* and *farauti 8*, respectively. Panel **C** shows nucleotide diversity (π, error bars show block-jackknife SE) and per-sample heterozygosity (Het.), grouped by population, across all taxa. Taxon (labelled across the top), with major geographic regions indicated in subheading. (**D)** and **(E)** are principal component analyses of *An. farauti s.s.,* and *An. hinesorum* and *An. oreios* respectively, with major geographic subgroupings labelled. Panel (**F)** shows the likely population graph topology recovered with *TreeMix* with 6 predicted migration edges (fig. S12). The graph is rooted with *An. punctulatus* (PUN). The strength of migration / admixture is indicated by the color of the arrow from yellow (light migration) to red (strong migration). No branch support is shown as nearly all are ≥ 90% (fig. S13). Populations are labelled by colour (see **B**). Major geographical regions of origin are labeled in grey to the right of the tips, with taxon labels in coloured italics. For clarity, population labels have been omitted from diversity statistics and *TreeMix* plot panels, fully labelled versions are in figs. S2 and S13.

A central aim of this study is to test whether the *An. farauti* complex shares evolutionary features that underlie the adaptive capacity of African and Asian anophelines: high standing genetic diversity structured by large segregating inversions, partially interbreeding cryptic taxa, and insecticide resistance driven by target-site and metabolic mechanisms. We further search directly for the genomic signatures of recent selection, testing whether documented shifts in vector behaviour, including changes in biting time and location following mid-20th-century DDT-based indoor residual spraying and 21st century ITN use (*14*, *38*, *39*), and the potential loss and re-emergence of human-biting in Western Province, Solomon Islands (*31*) – have a detectable genetic basis, or instead reflect behavioural plasticity acting without underlying selective sweeps.

We find that the *An. farauti* complex follows a distinctive evolutionary regime compared to other well-studied African and Asian *Anopheles* species complexes: highly structured, genetically depauperate populations shaped by repeated colonisation bottlenecks, with evidence of historical introgression from unsampled or extinct lineages. Selection at insecticide resistance loci is sparse and confined to a single population, and, with one possible exception, we recover no genetic signature underlying the behavioural variation that distinguishes these populations from one another. Our data indicate that the capacity for reported behavioural flexibility in this species complex need not be a recent, locally selected trait, and they describe a demographic context in which any adaptive response would be expected to be slow. The data generated in this study constitute a foundational genomic resource that will support surveillance of resistance evolution, the development of new control tools such as gene drive, and comparative study of speciation and adaptation across these species complexes.

### Climate and sea-level change fragmented a recently radiated complex

We generated whole genome sequences for 495 adult mosquitoes. From this genomic resource and after quality check and filtering, we analysed a genome-sequence dataset of 480 samples (median sequencing depth ∼30x, see supplementary materials), assigning them to 28 analysis populations on the basis of taxon, sampling locality, and genetic structuring (mean = 19, median=16, min=5, max=63) (see supplementary materials, tables S1 and S2, Fig. 1B–E). *An. farauti* s.s. and *An. hinesorum* were both sampled from Australia, New Guinea, Bougainville and the Solomon Archipelago, allowing comparisons between the two ecologically distinct taxa across a common geography, with additional *An. farauti s.s.* samples from Manus Island (politically part of PNG), and Vanuatu (the southeastern limit of its distribution) (Fig. 1B). *An. oreios* was sampled only from montane New Guinea (Fig. 1B).

Samples cluster strongly by species (fig. S1). *An. farauti* s.s. and *An. hinesorum*, the two most widely distributed and highly sampled members of the complex (Fig. 1B), share a common genetic structure across the region: each comprising three major genetic groups corresponding to Australia, New Guinea, and the island populations including the Solomon Archipelago (Solomon Islands together with Bougainville, an autonomous region in PNG). The *An. farauti s.s.* from Manus Island and Vanuatu also fall within the island group in *An. farauti s.s.* (Fig. 1D–E). In both taxa, low per-sample heterozygosity, per-population nucleotide diversity (π) and elevated individual long runs of homozygosity (ROH) – informative for effective population size and recent demographic history (*40*) – are most marked in the island populations, particularly Vanuatu and Bougainville respectively, followed by Australia, then New Guinea (Figs. 1C, 2A–C and fig. S2). This pattern is consistent across the single-taxa populations: *An. irenicus* and *An. punctulatus* sampled from the Solomon Archipelago show an elevated ROH burden (Fig. 1C and fig. S2). *An. oreios,* a sister species to *An. hinesorum* (*37*) sampled from the highlands of New Guinea (Fig. 1B), shows an elevated ROH burden compared with *An. hinesorum* from the lowlands of New Guinea (Fig. 2C). The parallel geography of structure and diversity loss across *An. farauti s.s.* and *An. hinesorum* is consistent with colonisation from a New Guinean centre of diversity, followed by sustained bottlenecks in island populations (*26*).

**Fig. 2.**
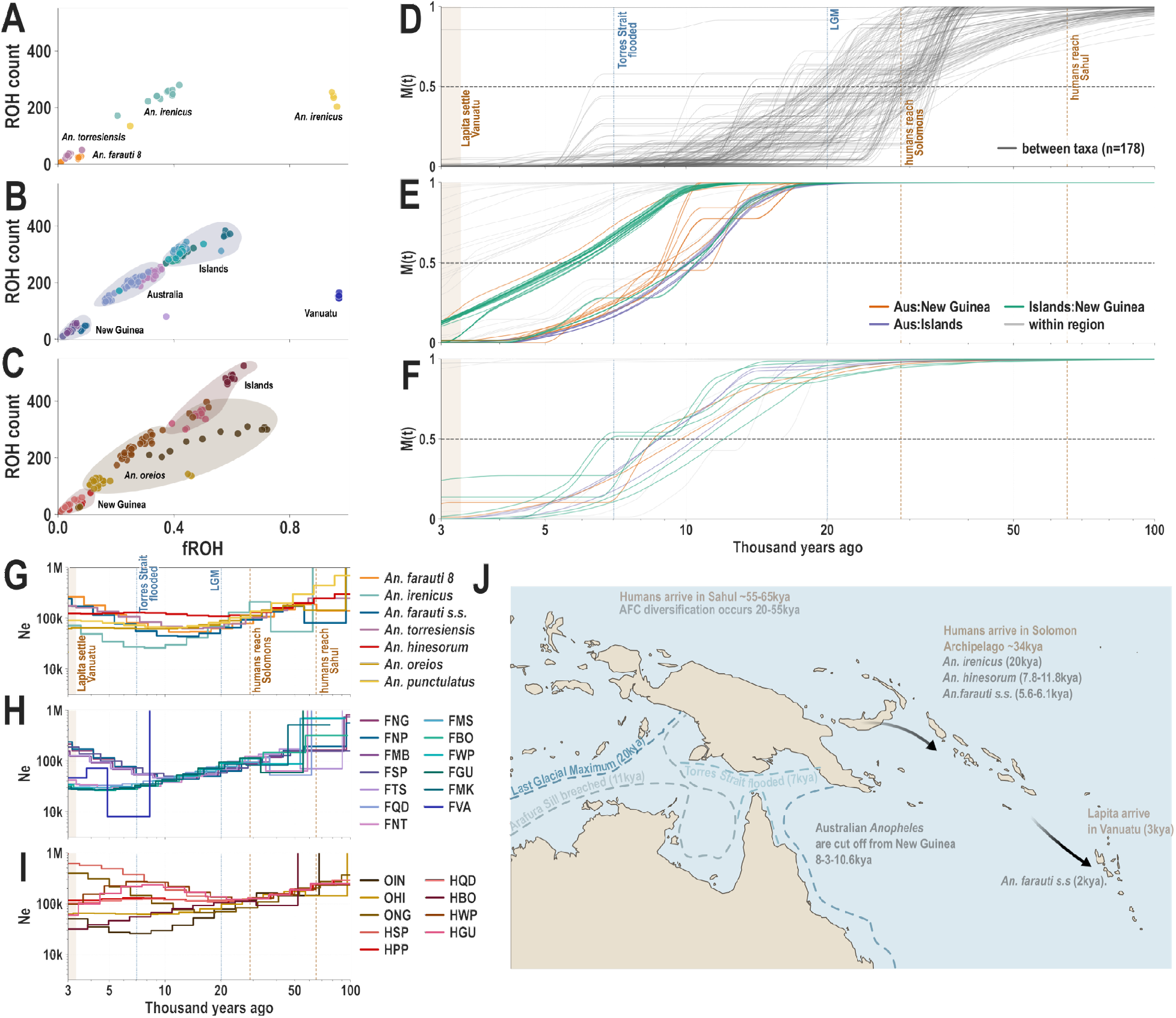
Population history and demography of the *An. farauti* complex. **(A to C)** Shows contemporary per-sample long runs of homozygosity (≥100kb) for the single-species populations (*An. punctulatus, irenicus, farauti 8* and *torresiensis*), *An. farauti s.s.*, and *An. hinesorum and An. oreios,* respectively. Taxon or major geographic regions are marked on plots. **D, E, and F** show cumulative migration probabilities (M(t), y axis), for all between-taxa population pairs, and within *An. farauti s.s*, and *An. hinesorum s.s* and *An. oreios*, respectively. M(t) is plotted over time (thousand years ago, x axes) at taxon-level population splits, and estimated with MSMC-IM. The dotted grey line shows M(t) of 0.5, the midpoint of population separation (*42*). **(G, H and I)** indicate taxon-level changes in effective population size (*Ne*), where the y axis indicates log(*Ne*), and the x axis, time (log10 years ago). Line colour indicates taxon. Note that taxon *Ne’s* for *An. farauti s.s, An. hinesorum,* and *An. oreios,* were computed from a single New Guinea population per taxon (see supplementary materials). Biogeographic events are shown as coloured lines on the plot, labelled above the top plots. For clarity, the M(t) lines in the all-taxon plots **D** and **E** are coloured depending on comparison level: between Australia, New Guinea, and island populations, or within-region. Note x axes across all plots is cut off at 3,000 years ago as per (*41*, *42*) (see **Methods**) as this is likely the limit of informative MSMC-IM resolution. (**J**) shows estimated sea levels from the Last Glacial Maximum ∼20kya to the breaching of the Arafura sill ∼10kya, to the inundation of the Torres Strait from 7kya - present, as well as colonisation events of humans and the AFC.

Across the complex in the island and Australian populations, inbreeding burden falls well above the values reported for major African vectors *An. gambiae* and *An. funestus* (*4*, *32*) (Figs. 1C, 2A–C), whereas the PNG populations are comparatively less inbred. *An. farauti s.s.* and *An. hinesorum,* differ markedly in how genetic variation is organised. Both show strong isolation-by-distance, consistent with strong barriers to gene flow imposed by common barriers to dispersal and island colonisation (fig. S3). *An. farauti* s.s. shows greater evidence of gene flow within each of Australia, mainland PNG, and the Solomon Archipelago, whereas *An. hinesorum* and *An. oreios* show more complex spatial structure at finer scales within each of these regions (figs. S4 and S5). Diversity is likewise distributed differently: in *An. farauti* s.s., π and heterozygosity are uniformly low across populations, with a minimum in Vanuatu where they approach zero, whereas *An. hinesorum* and *An. oreios* retain considerably higher and more variable diversity, particularly in mainland New Guinea (Fig. 1C and fig. S2). The result is a specialist species that is connected locally but depauperate everywhere, and a generalist one that is fragmented locally but retains more standing variation. Local PCA additionally reveals polymorphic inversions segregating in *An. hinesorum* and *An. oreios* (fig. S6), whilst the remaining taxa appear largely homosequential.

Our analyses suggest that this structure is the product of a rapid Pleistocene radiation. Reconstructing effective population size and cumulative migration rates through time (*41*, *42*) (see supplementary materials, Fig. 2D–I, figs. S7 to S11), we find that the *An. farauti* complex diversified rapidly in a single heterogeneous radiation between 20-55 thousand years ago (kya), with most taxa diverging in a 20-34 kya cluster (Fig. 2D, fig. S7). This divergence period was accompanied by a shared *Ne* decline around the time of the Last Glacial Maximum (LGM), ∼20kya, followed by a rebound and diversification of *Ne* trends, as the species-level divergences tapered off (Fig. 2G–I). The diversification resulted in an ecological spread spanning a coastal specialist (*An. farauti s.s*.), freshwater generalists (*An. hinesorum, An. farauti 8, An. torresiensis*), a highland specialist (*An. oreios*), and island zoophilics (*An. hinesorum* in the Solomon Archipelago and *An. irenicus* on Guadalcanal) (Fig. 1A).

The constituent taxa of the *An. farauti* complex then embarked on divergent evolutionary trajectories, shaped by their ecology. *An. farauti* s.s. has the lowest nucleotide diversity in the complex, distributed almost uniformly across its range (Fig. 1C and fig. S2). Its separation history is correspondingly cohesive: after a shared post-LGM *Ne* decline, two pulses divide the taxon: New Guinea from Australia at 8.8-10.6 kya, and New Guinea from the Solomon Archipelago at 5.6-6.1 kya, the latter separating Manus and all four Solomon Archipelago populations in a single synchronous event (Fig. 2E, figs. S7, S9, and S10). As a coastal species which utilises brackish water for larval habitats, this taxon plausibly experienced post-glacial sea-level change as a single, spatially coherent forcing (Fig. 2H): marine inundation of low-gradient coastal plains destroyed habitat in some regions while generating extensive new brackish water bodies in others, for example, in the Fly River delta and Sepik-Ramu lowlands (*43*, *44*), which we take to explain both the shared *Ne* increase in New Guinea populations between 10 and 5 kya and the tight clustering of separation dates within each pulse (Fig. 2H, figs. S7 and S8).

*An. hinesorum* and *An. oreios*, lowland and montane respectively, show the opposite pattern to *An. farauti s.s.*: higher and more variable diversity (Fig. 1C), more divergent demographic trajectories, and deeper, more staggered separations. *An. oreios* separates from *An. hinesorum* at ∼23 kya, coincident with the LGM - consistent with expansion of open montane habitat at peak cooling, when the New Guinea treeline was depressed by ∼1,200m (*45*), followed by fragmentation as that habitat contracted and moved upwards during Holocene warming, as observed in other montane taxa (*46*). Within New Guinea, *An. hinesorum* splits are older than their *An. farauti* s.s. counterparts (∼7.5-8.3 kya; Fig. 2E–F, figs. S7 and S8), and colonisation of the Solomon Archipelago is staggered across five millennia rather than synchronous: Bougainville at 11.8–12.8 kya, Guadalcanal at 8.3-8.8 kya, Western Province at 6.8–7.8 kya, with gene flow resuming among them only from 1.5-2 kya (Fig. 2E, figs. S7 and S10). Because these biome shifts were governed by local topography and rainfall rather than a single regional shoreline, the resulting separations are dispersed across a far wider window and lack the synchronous structure of *An. farauti* s.s. *An. irenicus* sits outside this pattern entirely. Restricted to Guadalcanal, it separates from its sister taxon *An. farauti* s.s. at ∼20 kya – more than threefold older than *An. farauti* s.s.’s own arrival on that island, indicating an independent and far earlier colonisation. It also carries the two deepest outlier splits in the radiation (×*torresiensis* 50.2 kya, ×*oreios* 55.2 kya; fig. S7) and is not equidistant from *An. hinesorum* despite its sister relationship to *An. farauti* s.s., and has low π and elevated ROH (Figs. 1C, 2A) all consistent with sustained isolation at small effective size on a single island.

Despite largely divergent demographic histories both *An. farauti s.s.* and *An. hinesorum* share a single event in common: the separation of New Guinea from Australia: *An. hinesorum* separates from Australia at 8.3–9.9 kya (figs. S7 and S9), closely tracking the equivalent split in *An. farauti* s.s. and the marine inundation of the Carpentaria region and Torres Strait between 10 and 6 kya (Fig. 2E,F,J).

Finally, we asked whether this history involved gene flow between taxa, as reported previously within *An. hinesorum* (*29*) and documented extensively in other major vector complexes (*47*, *48*). *TreeMix* (*49*), which fits a maximum-likelihood population tree from shared and private drift while adding migration edges that explain residual allele covariance, recovers a topology concordant with the genome-wide neighbour-joining tree (Fig. 1F, figs. S1, S12, and S13), with six supported migration edges (fig. S12) concentrated at deeper nodes. Resolving this with *Dsuite*’s *f*-branch statistic (*50*), which decomposes excess derived-allele sharing onto individual branches of the species tree across all 2,300 population trios, localises most of the signal to *An. irenicus* and *An. farauti* 8 and to basal branches (Fig. 1F and fig. S14). Because both methods are constructed to separate gene flow from incomplete lineage sorting, the most parsimonious source is one or more unsampled or extinct lineages closely related to the donor.

Several cross-taxon M(t) curves show extended plateaux rather than a clean monotonic decline (Fig. 2D), consistent with intermediate cross-coalescence maintained by this historical exchange. The most striking signal is in *An. hinesorum* from Western Province, Solomon Islands, where the presence of Papuan mitochondrial haplotypes was detected coincident with the emergence of human biting in a previously non-human biting population. We recover excess allele sharing between the two populations (fig. S14), suggesting a nuclear admixture event has taken place, though we are unable to localise this signal to a specific genomic region (fig. S15)

### Bottlenecked island *An. farauti* populations have purged deleterious variation

African *Anopheles* species are able to draw on deep mutational reserves of standing genetic variation, through strong positive selection at resistance loci, to resist chemical insecticide control (*51*). By contrast, the *An. farauti* complex shows a species-wide depletion of genetic variation relative to other *Anopheles* vectors for which comparable whole-genome data are available (*3*, *4*, *32*, *33*, *35*). This signal is pronounced in *An. farauti s.s*, and particularly in island *An. farauti s.s., An. hinesorum, An. irenicus* and *An. punctulatus,* and montane *An. oreios*, which show reduced nucleotide diversity (π) and contemporary effective population sizes (*Ne*), and a starkly increased ROH burden relative to populations from lowland New Guinea and Australia (Fig. 2D–E 3). Long term reductions in diversity and genetic bottlenecking as a consequence of island colonisation and population fragmentation may lead to the accumulation of deleterious mutations (genetic load), and a reduction in fitness as a consequence (inbreeding depression, genetic erosion), hampering the ability of the *An. farauti* complex to adapt to vector control.

We investigate this in more detail by estimating genetic load (see supplementary materials) across the *An. farauti* complex. Contrary to the expectation that the complex’s most bottlenecked taxon would carry the heaviest mutational burden, realised (homozygous) and masked (heterozygous) load at high-impact loss of function (LOF) and moderate-impact (missense) sites is lower in *An. farauti s.s.* than in *An. hinesorum* and *An. oreios* (Fig. 3A–B), with intermediate levels in *An*. *irenicus* and *An. farauti* 8 and *torresiensis* (Fig. 3A–B). Masked (heterozygous) load follows the same pattern (fig. S16). The taxa carrying the most deleterious variation are therefore those with the highest nucleotide diversity, heterozygosity and contemporary *Ne* (Figs. 1C, 2G–I) – the inland generalists – rather than the coastal specialist whose populations have most likely undergone repeated bottlenecks and persisted at small effective sizes. This is the signature expected where mutational input scales with *Ne* and where sustained small population size has allowed recessive deleterious variants to be exposed and removed through active purging. We find that island *An. farauti* populations show a deficit of homozygous high-impact mutations compared to their mainland counterparts (Fig. 3C), and compared to *An. hinesorum* (Fig. 3C), pointing to purging in these populations. By further finding that deleterious mutations are depleted within runs of homozygosity relative to synonymous sites, we find a plausible mechanism: recessive deleterious mutations entering ROH are exposed to selection.We find the opposite signature in Australian *An. farauti,* suggesting very inefficient purging of deleterious alleles in these populations. Purging in the *An. farauti* complex therefore appears to be concentrated where homozygosity is highest and effective size has been smallest for longest – the island populations of the coastal specialist – and is correspondingly weak in the more diverse inland generalists.

**Fig. 3.**
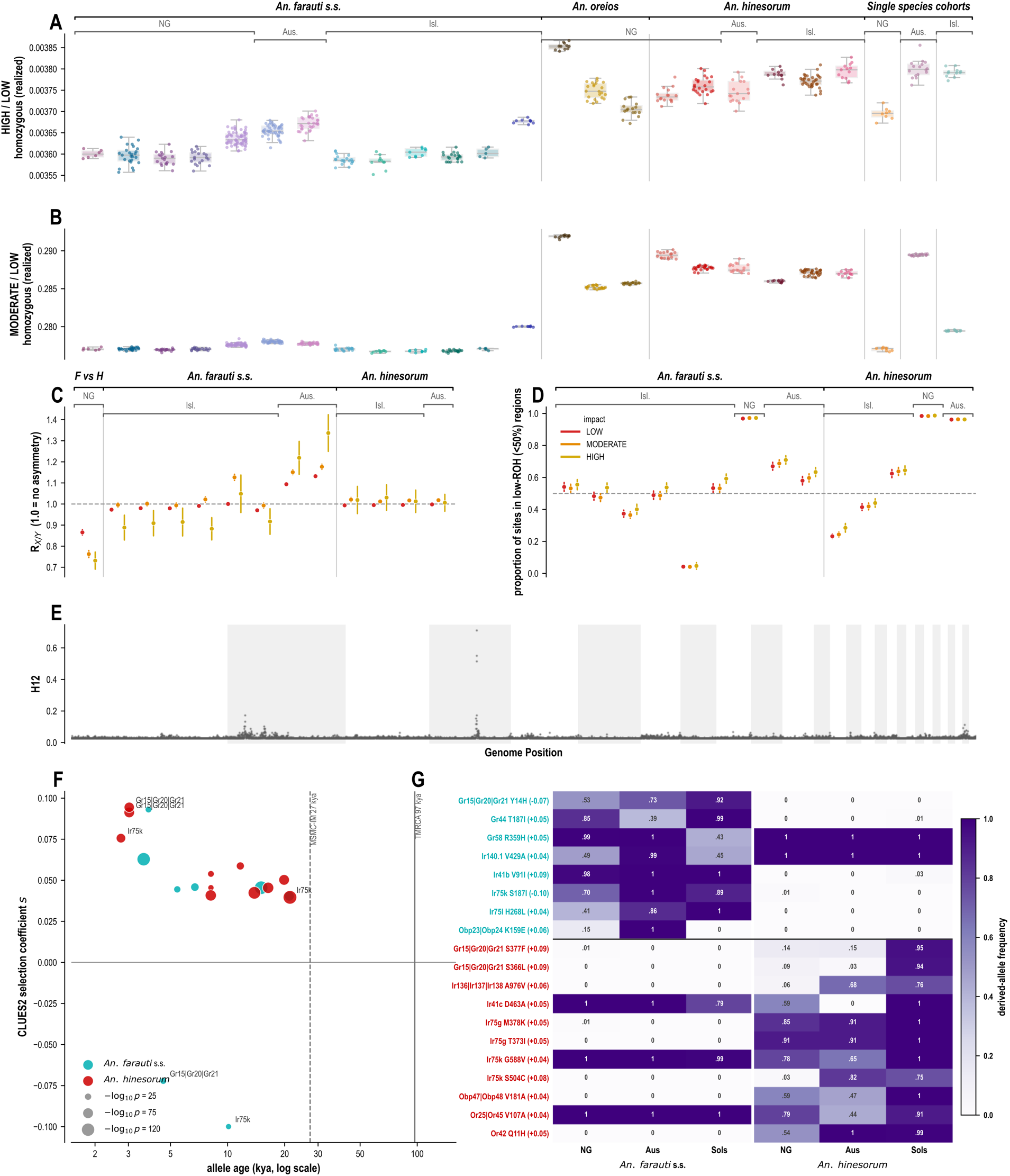
Adaptation and genetic load in the AFC. **(A and B)** show mutational load: per-sample derived-allele burden at HIGH- (**A**) and MODERATE (**B**)-impact sites, expressed as a ratio to synonymous (LOW-impact) sites to control for each taxon’s own demographic history. Boxes show population medians; points are individuals. **(C)** shows *R_x|y_*: relative deleterious load between each island population and its own taxon’s New Guinea mainland baseline (block-jackknifed 95% CI; dashed line = 1, no difference). Island *An. farauti s.s.* sit below 1, consistent with purging, and Australian *An. farauti* far above one, showing an excess of deleterious mutations. Island *An. hinesorum* shows no difference. **(D)** ROH-purging: the proportion of segregating deleterious sites falling in low-runs-of-homozygosity regions (dashed line = 0.5). HIGH/MODERATE sites skew toward low-ROH in island *An. farauti* s.s. populations, showing the purging mechanism, inbreeding exposing recessive deleterious alleles to selection. No difference is evident in *An. farauti s.s.* from Vanuatu, as most of the genome is in ROH, or the rest of the island populations in the complex.. In panels **A-E**, taxon and geographic region are labelled, but for clarity, individual population labels have been hidden and can be found at figs. S16 and S19. **(E)** Is a genome-wide selection scan using the H12 statistic, of *An. farauti s.s.* in Guadalcanal. Y axis denotes H12, x axis denotes genomic position. Alternating shading bands indicate separate contigs. The only apparent selective sweep, at the *Gste2* locus, is labelled. **(F)** shows CLUES2 selection coefficients, and mutational ages, at 19 missense SNPs in 206 chemosensory associated genes in *An. farauti s.s.* and *An. hinesorum*. Y axis denotes selection coefficient (where positive values indicate positive selection, and vice versa), x axis denotes mutational age in years. Dotted line indicates the MSMC-IM inferred divergence time between *An. farauti s.s.* and *An. hinesorum,* the solid line, the same for the AFC and *An. punctulatus.* Text labels show variants in genes under selection in both taxa (**G**) shows the derived allele frequencies for the 19 significant missense SNPs (x axis, labelled with their respective selection coefficients) across the three major geographic regions in *An. farauti s.s.* and *An. hinesorum*.

### Recent selection is rare and species-specific

The demographic histories reconstructed above bear on the capacity for adaptation. Purging shows that selection acts efficiently against strongly deleterious recessive variants in these populations, but small effective size and depleted standing variation limit the supply of variants on which positive selection could act. We therefore asked whether these populations show any signature of recent positive selection, and if so, at which loci.

Extensive indoor residual spraying (IRS) campaigns with DDT were carried out in PNG and Solomon Islands (*14*, *38*) in the mid-20th century, followed by widespread ITN and IRS campaigns in the 21st century. In African and South Asian *Anopheles,* these efforts have led to the development of widespread physiological resistance, which has itself left strong, repeatable signatures of positive selection in vector genomes (*32*, *35*, *51*, *52*). However, phenotypic resistance in the Pacific seems rare and highly localised (*11*, *12*), with sub-optimal control response instead attributed to widespread outdoor and early biting (*10*, *14*, *53*), and ITN durability issues (*13*). Accordingly, genome-wide selection scans of *An. farauti s.s., An. hinesorum* and *An. oreios* samples collected across PNG and Australia by *Operation Anopheles,* prior to mass ITN rollout, find no evidence of selective sweeps at resistance loci (figs. S17 and S18). Similarly, in contemporary samples collected post ITN-rollout (2018-present) from Solomon Islands and a single location in PNG (Madang, FNC), we only see a single localised sweep at the *Glutathione-s-transferase epsilon subfamily 2* (*Gste2*) cluster in Guadalcanal *An. farauti s.s.* (Fig. 3E).

While physiological resistance seems largely absent, the *An. farauti* complex has long been studied as a system in which control failure is mediated behaviourally rather than physiologically (*9*, *10*) and the traits concerned – host preference, biting time, and biting location – may vary widely within and between taxa (*6*, *16*, *26*, *53*). Unlike other major vectors, it has undergone multiple transitions from anthropophily to zoophily (*6*, *26*). Our previous work identified taxon- and population-specific divergences in candidate genes belonging to several gene families involved in insect chemosensation, including gustatory proteins (*Gp*s), olfactory receptors (*Or*s), ionotropic receptors (*Ir*s) (*19*) and odorant-binding proteins (*Obp*s) (*30*), but we could not resolve which populations selection acted on, or when (*30*). Revisiting this dataset, screening 206 candidate chemosensation-associated genes (see supplementary materials and tables S3 and S4) for taxon-specific selection coefficients with Relate (*54*), identified 19 significant (P < 5 × 10^−8^) missense sites in total across 14 unique transcript IDs in *An. hinesorum* (n=11) and *An. farauti s.s.* (n=8), with mutations in the *Ir* (n=8)*, Gr* (n=5)*, Or* (n=3)*, and Obp* (n=3) families (Fig. 3F–G). Of these, 17 were significant for positive selection, and two for negative selection. None of the mutations were shared between species, and all of their origins post-date the *An. farauti-An. hinesorum* split (table S4). Two transcripts, corresponding to *Gr15|20|21* and *Ir75k* have positively selected mutations in *An. hinesorum,* and are negatively selected in *An. farauti* (Fig. 3F). Whilst our analysis is underpowered to detect selection at sub-taxon level, the two variants in *Gr15|20|21* are present at very high frequencies in *An. hinesorum* from Solomon Archipelago, and very low frequencies elsewhere (Fig. 3G), with an approximate allele age of 3kya (Fig. 3F), post-dating the colonisation of the islands by *An. hinesorum.* None of the genes identified here were among the candidates nominated by our previous divergence-based analysis of nine genomes (*30*), suggesting that recent selection has acted on chemosensory genes other than those bearing the signature of long-term divergence.

### A distinct evolutionary regime for malaria mosquitoes in the southwest Pacific

Malaria vectors in the southwest Pacific have followed an evolutionary path that looks vastly different to that in Africa. The radiation of the *An. farauti* complex overlaps with the peopling of Sahul (*55–57*). The presence of hominins has also been linked to the diversification of anophelines in Southeast Asia (*58*), as well as speciation events in *An. gambiae* (*59*) and *An. funestus* (*60*). Post-glacial sea-level rise then severed New Guinea from Australia and separated mosquito populations with it (63), with island colonisation proceeding through a series of founder events. Its two most widespread members responded to that shared history in opposite ways. The coastal specialist *An. farauti* s.s. experienced sea-level change as a single coherent forcing and became uniformly depauperate. The inland generalists *An. hinesorum* and *An. oreios* tracked more heterogeneous shifts in terrestrial habitat and retained more variation. The result is a set of isolated, inbred populations lacking almost every feature behind rapid resistance evolution in African vectors: large effective size, abundant standing variation, connectivity and adaptive introgression (*3*, *4*, *32*), (*61*, *62*). That this contrast has gone unrecognised reflects how little is known about the region’s vector biology at the genomic level. Near Oceania offers a dated sequence of sea-level, climatic and colonisation events against which molecular evolution can be calibrated. The history of its human populations has been reconstructed in detail (*63*) but the history of its insects, including the vectors that shaped its disease burden, has not.

Chemical control has seen extensive use in the region. DDT was sprayed in parts of PNG from the 1960s and across Solomon Islands in the 1970s, where it is widely credited with shifting *An. farauti* toward early-evening and outdoor biting (*14*), although that evidence rests on a handful of sites and predates molecular species identification. ITN coverage in PNG has since been broadly comparable to that in sub-Saharan Africa (*5*, *5*, *64*, *65*). Our sampling spans this history, from archived pre-ITN collections across PNG and Australia to contemporary material from Solomon Islands and Madang. Yet across three decades we find a single resistance sweep, at *Gste2* on Guadalcanal, where both isolation and exposure history are most extreme. This is consistent with phenotypic surveillance, although that surveillance has been sparse outside of PNG (*11*, *66*). Nor do we find selective sweeps or structure pointing to a heritable basis for the region’s divergent biting behaviour. That includes zoophilic *An. hinesorum* in Solomon Islands versus its anthropophilic relatives in New Guinea, a contrast with *An. arabiensis*, whose host choice tracks the 3Ra inversion (*17*, *18*). The one exception may be excess allele sharing between *An. hinesorum* of the Papuan Peninsula and Western Province, Solomon Islands. It is consistent with the mitochondrial introgression reported previously (*29*) and with the emergence of human biting there (*31*), but it does not relate to any genomic region.

The simplest explanation is that selection has been weak, for several reasons. Nets distributed in PNG have shown reduced bioefficacy (*13*). Early, outdoor and zoophilic biting behaviour, which is widespread across the complex and apparently ancestral rather than derived (*6*, *15*, *16*), limits contact with treated surfaces. Spraying may also have removed the more endophilic taxa or individuals rather than selecting within them, as is suggested by the species turnover recorded after DDT campaigns (*14*, *67*). Where a resistance allele does arise, the strong barriers to gene flow we document could slow its spread. Behaviour that blunts indoor control may therefore have shielded these vectors from the selection imposed by control. The genetic controls of biting behaviour are likely highly polygenic. Our study is not powered to detect them, but they provide data as a platform for the association studies that could.

Extensive inbreeding also has not been a liability. Island *An. farauti* s.s. are among the most inbred populations in the complex – with Vanuatu almost completely homozygous – yet they carry fewer loss-of-function variants than their New Guinea progenitor population (Fig. 3C), consistent with recessive deleterious alleles being exposed and purged as autozygosity rose (Fig. 3D). The more diverse inland generalists carry more. The complex thus contains distinct adaptive regimes: a coastal specialist that has been small for long enough to be scoured of much of its deleterious variation, and inland taxa that more closely resemble vectors elsewhere.

Throughout, it persists, abundant and ecologically diverse, under demographic conditions usually expected to erode fitness. *An. stephensi* offers a parallel: a severely bottlenecked invader that has spread rapidly across the Horn of Africa carrying resistance acquired before it arrived (*35*). In both cases, as in the urban *Culex pipiens* form *molestus* (*68*) success rests on traits already in place rather than on new adaptation. What such populations may lack is the capacity to respond to fresh selection. Low standing variation, small effective size and limited gene flow should slow both the emergence and the spread of resistance, a prediction this complex is positioned to test as control intensifies.

That a complex this depleted and this inbred remains capable of sustaining intense transmission has implications beyond malaria, for how persistence under severe demographic constraint is understood. It also bears directly on control: as new interventions reach the region, from spatial emanators (*69*) to the prospect of gene drive, for which low diversity at target sites is an advantage, a genomic baseline will be needed to anticipate how vectors respond. This work provides the first such foundation for malaria vectors in the southwest Pacific, with the data openly available to support these efforts.

## Acknowledgments

We wish to thank community members and leaders who consented to sample collections, all sampling teams and ADF staff and helicopter pilots involved in mosquito collection efforts. We are grateful to Etienne Patin and Mathilde Andre for their discussion of human population history in the region. This research relied on the Institut Français de Bioinformatique HPC Core Cluster facility (ANR-- 11-- INBS-- 0013) and the ISO--9001 certified IRD i-Trop HPC (member of the South Green Platform) at IRD Montpellier, and the Research Computing Centre at the University of Queensland for their provision and maintenance of the excellent Bunya supercomputer. Finally, we thank Cynthia Riginos, David Weetman and Martin Donnelly for their insightful feedback.

The opinions expressed here are those of the authors and do not necessarily reflect those of the Australian Defence Force and/or extant Defence Force Policy.

The authors acknowledge the use of Claude Opus 5 (Anthropic, San Francisco) in the creation of this work. All analyses and decisions were designed by the authors, who parametrised initial analysis code and workflows for each analysis in the manuscript. Claude was then used to troubleshoot, optimise job batching, and tidy code for publication. All conceptualisation, analysis, and interpretation were performed by humans. All code has been checked for consistency and correctness by the authors. The manuscript was entirely written and reviewed by humans. The authors take full responsibility for the code-base and manuscript.

## Author contributions

Conceptualization: TPWD, LA, MCF, NWB Data curation: TPWD, LA

Formal analysis: TPWD, LoT, MCF Funding acquisition: NWB

Investigation: TPWD, LA, FB, LoT, WKC, TR, DW, MJB, RV, MK, LiT, SL, PJ, AFvdH, GD, AWS, SPF, TRB, HB, SK, RDC, MCF, NWB

Methodology: TPWD, MCF Project administration: NWB

Resources: TPWD, LA, FB, WKC, TR, DW, MJB, RV, MK, LiT, SL, PJ, AFvdH, GD, AWS, SPF, TRB, HB, SK, RDC

Software: TPWD, MCF Supervision: MCF, NWB Validation: TPWD

Visualization: TPWD, LoT

Writing – original draft: TPWD, MCF, NWB

Writing – review & editing: TPWD, LA, FB, LoT, WKC, TR, RV, MK, LiT, SL, PJ, AFvdH, GD, SPF, TRB, HB, SK, RDC, MCF, NWB

## Data, code, and materials availability

Code is available at https://github.com/tristanpwdennis/afc-evolution-paper. Raw reads are deposited at the European Nucleotide Archive (ENA) under project number PRJEB130419. SNP genotype data will be made available through public repositories upon manuscript acceptance.

## Supplementary Materials

Materials and Methods

Figs. S1 to S21

Tables S1 to S4 References (*71–117*)

