## Supplementary material for "Population genomics of the *Anopheles farauti* complex reveals a distinct regime of malaria vector evolution in the southwest Pacific": Methods and Supplementary Material

**The PDF file includes:**

Materials and Methods
Figs. S1 to S21
Tables S1 to S4

### Materials and Methods

Sample collection information and population delineation

For a full breakdown of sample collection dates and exact locations, please see table S1.

Papua New Guinea

In mainland PNG, samples were collected between 1992-1998 using human-landing catches, $CO_2$-baited traps, light-traps, and larval dips, as described previously (*23*).

On Manus and Bougainville, samples were collected in 2019 and between 1998-1999, respectively.

Contemporary samples were collected from Papua New Guinea in Madang Province in larval surveys and human landing catches. Owing to compute constraints, these were processed identically but analysed separately from the rest of the callset (see below), and are labelled as population FNC.

Solomon Islands

On Guadalcanal, samples were collected in 1997, then in additional trips in 2015 and 2018 (*30*). In Western Province, samples were collected in 2016 (*30*) and 2023 .

Australia

Australian samples were collected between 1992-1998 using human-landing catches, $CO_2$-baited traps, light-traps, and larval dips, as described previously (*24*). Contemporary Australian samples were collected using the following methods: (i) a novel mosquito catch barrier (MCB) system, (ii) $CO_2$-baited Fan Box Traps (FBTs), and (iii) human landing catches (HLCs). FBT and HLC collections were conducted according to previously described protocols (*71, 72*)

Vanuatu

Vanuatu samples were collected using human-landing catches in 2008.

Indonesia

Samples were collected using human-landing catches in West New Guinea (then Irian Jaya), Indonesia between 1996-1999 (*38*).

DNA Extraction and species PCR

Mosquito DNA was extracted using a salt-based extraction technique (*73*). We used restriction fragment length polymorphism (RFLP)-based assay to discriminate within members of the *An. farauti* complex (*74*), and assign species status. 520 samples were sent for sequencing (Illumina 2x150BP paired-end reads) at a commercial provider. Of these, 515 passed library QC and were successfully sequenced.

Bioinformatics

Versions of the software used in this study are given in the GitHub repository.

Read mapping and variant calling

Throughout this study, we used the Anop_fara_FAR1_V2 *An. farauti* assembly reference genome (*75*), downloaded from Vectorbase Release 71(*76*). We followed a standard pipeline for variant discovery, incorporating BWA-MEM (*77*), GATK's MarkDuplicates (*78*) and GATK

HaplotypeCaller (*79*) to produce individual .gvcf files. As the *An. farauti* reference assembly is not assembled into chromosomes, we limited alignment and variant discovery to the 46 largest contigs, representing >90% of the assembly. Individual .gvcf files were jointly genotyped with GATK's GenomicsDBImport and GenotypeGVCFs commands in parallel across all called contigs. Per-contig all-sample .vcf files were converted in zarr datastores (*80*).

<u>Sample quality control</u>

We performed sample quality control (QC) on our initial callset of 515 individuals. We computed the per-individual median and mean depths-of-coverage, and the fraction of the genome covered by at least a single read. We excluded samples with a median depth-of-coverage of < 8, and a fraction covered < 0.8. This resulted in 11 samples being excluded from the callset. Additionally, we computed principal component analysis (PCA) and neighbour joining-trees (NJT) from 100,000 randomly selected variants from the largest contig (KI109040) to check for possible contamination or misspecified species assignment from the initial RFLP/PCR assay (*74*). We excluded an additional 9 samples based on positioning in NJT/PCA on extremely distal branches / locations, indicating possible contamination or misspecification. Furthermore, we identified 6 samples that appeared to be misspecified based on their speciation RFLP/PCR assay and positioning in the NJT/PCA – e.g. samples marked as *An. farauti sensu stricto* grouping with *An. hinesorum*. We updated their taxon label accordingly. We updated 40 taxon labels, including 31 samples marked as *An. hinesorum* that grouped with *An. oreios.*The final qc-passing set contained 495 samples. For 15 of these, we were not able to recover accurate geo-location information. These samples were included in phasing ARG inference to bolster sample sizes, but were excluded from any population- or sampling-location stratified analysis. This left a core analysis set of 480 samples (table S3).

<u>Site quality control</u>

We generated a static site filter by computing sitewise summary statistics from all samples (mean mapping quality (MQ), genotype quality (GQ) and depth (DP) across all samples retained by sample QC. We marked sites as PASS (True) where:

- Mean MQ > 50
- Mean GQ > 60
- No more than one individual with a DP of 0
- < 10% of individuals have at least half or less the modal coverage
- < 5% or more individuals have at least twice the modal coverage
- The site was *not* an INDEL or COMPLEX site

This excluded approximately 20% of total sites from the callset.

<u>Statistical phasing</u>

We used Beagle v5.5 (*81*) to phase the whole callset, with a window size of 10cM, and an overlap of 1cM. In the absence of a genetic map for *An. farauti*, we assumed a constant recombination rate of 1cM/Mb.

<u>Population delineation</u>

Samples were grouped into 28 distinct analysis populations based on observed patterns of genetic structuring, or by geographic location. See Fig. 1B and fig. S1, and table S2 for more information.

The 30 additional contemporary samples from Madang, Papua New Guinea, received after the main callset was generated, were processed identically but genotyped and phased separately owing to compute constraints. These samples were used only for the G123 contemporary selection scan and are excluded from all other analyses. Comparisons between this population and the archived northern PNG population were restricted to sites passing quality filters in both callsets.

Population Genetic Analysis

Population genetic data analysis and plotting were largely performed using the scikit-allel (*82*), numpy (*83*), pandas (*84*), sgkit (*85*), tskit (*86, 87*), seaborn (*88*), plotly (*89*), and xarray (*90*) libraries, as well as code adapted from the malariagen_data API (*91*).

Neighbour-joining tree analyses

We inferred neighbour-joining trees from 100,000 randomly selected SNPs from the largest inversion-free contig KI109040. City-block distances were inferred between all pairs of sites in all individuals. The resulting distance matrix was used to infer a neighbour-joining tree in the anjl (*92*) library, which implements the neighbour-joining algorithm of Saitou and Nei (*93*).

$F_{st}$

We computed the average genomewide $F_{st}$ between all chromosomes, using allele counts from all population pairs, in *scikit-allel* using Hudson's $F_{st}$ estimator, which is more robust to unequal sample sizes (*94*).

Isolation-by-distance

To estimate isolation by distance, we estimated the pairwise geographic distance between every site for which we computed $F_{st}$ (above). Within *An. farauti ss* and *An. hinesorum,* we regressed Haversine distance against linearised $F_{st}$: $1\text{-}1/F_{st}$. See fig. S3 for model statistics.

fEEMS

We inferred landscape genetic structure in the AFC using *fEEMS* (*95*), a software library for inferring 'migration surfaces'. 100,000 SNPs were randomly selected genomewide and used as input for *fEEMS*. Triangular grids, of spacing 75km, spanning the analysis region (the Southwest Pacific) were created using the *ddgridR* (*96*) library. The resulting grid, dataframe of sampling locations, and *PLINK* format SNPs, were used as input for *fEEMS*. Leave-one-out cross-validation was performed as per the *fEEMS* user manual, and the grids with the lowest cross-validation error were selected for plotting.

Thetas

We computed nucleotide diversity ($\pi$) and Tajima's $D$ in 10kb windows across the genome using *scikit-allel*, which, when given the chromosome start and end, calculates $\pi$ using all sites (including uncalled sites) as a denominator (*97*). 95% confidence intervals were calculated with block-jackknife resampling.

ROH

We inferred per-individual and per-chromosome runs of homozygosity (ROH) using *scikit-allel*, which uses a hidden markov model (HMM) to infer ROH. ROH were inferred from accessible SNPs genome-wide with a HMM (*4, 82*), and filtered to include ROH greater than 100,000bp in length (*4, 82*). Output was combined and aggregated to provide per-individual genome-wide estimates of the number of ROH in an individual genome, and the fraction of the genome in ROH. We filtered ROH to include only segments longer than 100kb (*4, 98*), to include only those putatively a result of recent inbreeding.

<u>Local PCA – inversion detection</u>

Population structure may vary across the genome (*99*). In the case of mosquitoes, heterogeneity may arise as a consequence of chromosomal inversions, which can be detected by performing PCA in windows across the genome (*33, 99*). We performed local PCA in windows of 50kb across all contigs using winPCA (*100*).

<u>TreeMix Analysis</u>

We investigated the relationships between the different populations with *TreeMix* 1.13 (*50*). This creates a maximum-likelihood population tree based on allele frequency correlations between populations. Reticulations or migration edges (m) are added between branches between populations that share ancestry through admixture or geneflow. We used 2,136,808 high-quality physically unlinked biallelic SNPs. This data set was obtained by filtering genotypes for quality (GQ) higher than 30, then keeping only SNPs with less than 5% missing data, and no missing data at the outgroup (*An. punctulatus*), and removing singleton mutations. Then physically unlinked SNPs are selected using LD-pruning with PLINK v1.90b7.2 (*101*) (option –indep-pairwise 500 - 250 - 0.1). The TreeMix analysis was run in three steps. First, we estimated the number of *m* that best fitted the data by running *TreeMix* 10 times for each *m* value, with *m* ranging from 1 to 10. The optimal *m* value was estimated using the Evanno method using the OptM R--package (*102*). Second, a consensus tree with bootstrap node support was obtained by running TreeMix 100 times using the optimal *m* value (m = 6), the --bootstrap option with resampling blocks of 100 SNPs (-- k 100), generating the consensus tree using PHYLIP *consense* tool (*103*). Finally, the third step runs a final TreeMix analysis using the consensus tree as a guide (--tf option) and all the data, with the best m value (*m* = 6). These steps were run as part of a modified version of the BITE pipeline and the associated R scripts (*104*). In each step, *An. punctulatus* was used as an outgroup to root the trees.

<u>f-branch analysis</u>

We used *f-branch* analysis in *Dsuite* (*51*) to examine in more detail potential gene flow and introgression events across the AFC. Taking the best-fit topology from the TreeMix analysis (*m*= 6), *Dsuite* assigns correlated D-statistic / f-4 ratio (*105–107*) introgression signals to specific branches of the AFC tree, resolving whether admixture signal sits on terminal branches (recent) or on internal / ancestral branches (potential older admixture or incomplete lineage sorting). We used per-contig phased multisample VCFs (see *Phasing*) as input, computing *D* and f4-ratio statistics per population trios. We merged the 46 per-contig outputs into a genomewide trio-statistic, rearranging them according to the input species tree (see above). We then used *f-branch,* which uses the trees branching structure to decompose correlated trio signals into per-branch f-branch ($f_b$) statistics, plotted as a heatmap.

Demographic history and split-time inference (MSMC2 / MSMC-IM)

We inferred within-population effective population size histories, Ne(t) and between-population split-times using the multiple sequentially Markovian coalescent (MSMC-2 v2.1.4) (*42*), together with the isolation-migration (-IM) extension (*43*).

In the absence of a genome-wide callability mask, we used the filter mask (see above) to estimate callability over phased sites (see above for phasing). Switch-error noise is therefore expected to inflate coalescence-rate estimates in very recent time-scales) but will not bias the absolute order of events.

Per-sample single-sample VCFs were generated per-contig and combined into multi-sample multihetstep files with multihetstep.py in msmc-tools (*108*). Individuals were assigned into two groups: taxon and population (tables S1 and S2). Because MSMC2 scales poorly with sample number, each group was represented by a fixed subset of 4 diploid individuals (8 haplotypes) for within-population runs, and the first two individuals that group for cross-population runs (4+4 haplotypes), drawn at random with a fixed number seed. For the three taxa with internal population structure: *An. farauti ss, An. hinesorum* and *An. oreios,* single populations (FSP, HPP, HOI) were used to represent each taxon, as the PNG populations represent the original centers of diversity for these taxa. Estimates of Ne(t) and relative cross-coalescence rate (RCCR) of these runs were consistent with the between taxon population level runs (Fig. 2 and figs. S7 to S11).

Within-population inference

For each group, MSMC2 was run on the 8 within-group haplotypes across all 46 phased scaffolds jointly (-I 0,1,2,3,4,5,6,7), using the default 32-segment time-pattern (-p 1*2+25*1+1*2+1*3), producing per-group coalescence-rate estimates (lambda00) and a single scaled cross-coalescence output per group.

Cross-population inference

For each of the taxon-pair and population-pair comparisons, MSMC2 was run with -s (skipAmbiguous) across all 16 between-group haplotype pairs (4 haplotypes/side; -I 0-4,0-5,...,3-7) using the identical time pattern, to estimate the cross-coalescence rate (lambda01). Within- and cross-rates for each pair were combined into a single joint rate table with combineCrossCoal.py (msmc-tools).

MSMC-IM fitting

The combined lambda00/lambda01/lambda11 rate table for each pair was fit with MSMC-IM (MSMC_IM.py), specifying the same 32-segment time pattern and a per-generation mutation rate of mu = $1 \times 10^{-8}$ site$^{-1}$ generation$^{-1}$ (matching scaling used elsewhere in this study), to obtain: (i) the relative cross-coalescence rate through time, rCCR(t) = 2*lambda01 / (lambda00 + lambda11); (ii) a symmetric isolation-migration model fit giving time-varying migration rate m(t) and cumulative probability of having split by time t, M(t); and (iii) population-specific scaled effective sizes under the IM model. Rates and times were converted to biological units using mu = $1 \times 10^{-8}$ site$^{-1}$ generation$^{-1}$ and a generation time of 11 generations/year (Ne = 1/(2*mu*lambda); t_years = t_scaled/mu/11) as per (*4*). Split times were taken as the point at which rCCR(t), read backward in time, first crosses 0.5 (rCCR50 kya), a split-time heuristic for MSMC2/MSMC-IM cross-coalescence curves, where rCCR = 1 indicates a single ancestral population and rCCR = 0.5 marks the point of population separation (*109*).

Ancestral recombination graph (ARG) inference.

We inferred ancestral recombination graphs across our sample set using Relate (*55*), on the phased callset above minus the six *An. punctulatus* individuals, which were used to polarise the data. We used a flat, genomewide recombination rate of 1.3 cM/Mb, to infer a pooled ARG per-contig, with a mutation rate of $1x10^{-8}$ per site, per generation (*4*), producing per-contig .anc and .mut files. These were then converted into tskit (*110*) format for further analysis using Relate's ConvertToTreeSequence module.

Variant functional annotation

Biallelic SNPs from coding sequences were extracted from the callset based on canonical mRNA transcripts from the VectorBase reference genome annotation (*75, 76*). Functional consequences were annotated per canonical mRNA transcript using a custom script (veff.py, see GitHub repository) from the malariagen_data Python API (*91*), assigning an effect and impact class (e.g. synonymous/LOW, missense/MODERATE and loss-of-function/HIGH) to each SNP. Where a site fell within more than one transcript, the highest impact class was retained. Genotypes were repolarised using the *An. punctulatus* outgroup. MODIFIER-class sites (intronic/UTR) were excluded from downstream analyses.

Per sample mutational load

For each sample, derived-allele genotypes were tallied per-impact class separately for heterozygous (1 copy, masked load) and homozygous (2 copies, realised load), and summed across all annotated contigs. Load was expressed as the ratio of HIGH/LOW and MODERATE/LOW consequence mutations, using the LOW as an internal neutral baseline to account for differences in *Ne* and drift between taxa and populations.

Relative deleterious load between groups - $R_{xy}$

Following previous work (*111*), we computed per-population $R_{xy}$, which compares the accumulation of derived deleterious mutations between populations. If selection has been equally effective and mutation rates constant since population split, $R_{xy}$ is expected to equal 1. In practice, founder effects and genetic drift shift the neutral (synonymous, LOW) classes $R_{xy}$ away from 1 as well, so we treat each population pair's own synonymous-class $R_{xy}$ as a null and interpret deleterious classes (missense/MODERATE, LOF/HIGH) relative to that baseline. Briefly, Rxy was computed as before (*111*), summed over segregating sites in a given impact class, comparing a focal group (X) against a reference group (Y). Confidence was assessed by block jackknife (100 contiguous, ~equal-SNP genome-order blocks across contigs. Comparisons were structured as island or Australian population vs. that taxon's pooled New Guinea (PNG-only) populations, and farauti_ss-PNG vs. hinesorum-PNG.

Purging via homozygous exposure - ROH test

We computed the fraction of realised mutational load inside ROH to detect possible purging. In populations where genetic purging through inbreeding has occurred, the homozygous mutation load should be smaller than expected given estimates of inbreeding. This is because purifying selection acts to reduce the frequency of deleterious recessive homozygotes, removing two deleterious alleles from the population per locus. If inbreeding-driven purging has occurred, deleterious alleles exposed as homozygotes inside ROH are preferentially removed by selection, so what remains segregating today should be depleted in high-ROH regions and enriched in low-ROH regions relative to neutral sites. For each population group, every segregating site (per impact class) was classified by the fraction of that group's individuals for whom the site falls inside a run of homozygosity (ROH; per-sample ROH calls from data/roh/), thresholded at 50% into low-ROH (<50% of individuals) vs. high-ROH (>=50%) sites. Then, we computed the proportion of segregating sites falling in low ROH regions separately for each impact class (LOW/MODERATE/HIGH) within each group. Groups were defined identically to the $R_{xy}$ analysis: per-taxon island populations plus pooled New Guinea (PNG-only) and Australia-by-state (QLD/NT) reference groups. Standard error estimates were computed using block-jackknife.

Genome-wide selection scans

We performed genome-wide selection scans (GWSS) for evidence of recent selection by using Garud's G123-statistics (*112*). G123 was computed in windows of 1000 haplotypes, aggregated and plotted by population. We stratified our populations into pre- and post- ITN distribution for plotting.

Chemosensory gene curation

We built the candidate chemosensory gene set from the 210 genes we previously manually annotated in the *An. farauti* complex (*31*): 54 odorant receptors (Or), 68 odorant-binding proteins (Obp), 38 ionotropic receptors (Ir) and 50 gustatory receptors (Gr). Extracted coding sequences were searched against the *An. gambiae* PEST (*113*) annotated proteome (VectorBase release 68) with BLASTX (*114*). We took the top hit as the orthologue and compared it with the ID implied by the published gene name. The name-based ID came from a lookup of the gene symbol against all 13,845 *An. gambiae* PEST gene records in VectorBase. Sequences were available for 208 of the 210 genes; *Or65* and *Ir40c* have no deposited sequence and were not assigned. The BLASTX top hit matched the name-based ID for 153 genes. For 28 genes a name-based ID could not be found and the ID was taken from sequence identity alone. Another 12 were off by one or two gene numbers, which we attribute to an older *An. gambiae* annotation. Three (*Ir136*, *Ir139*, *Ir140*.1) mapped to newer IDs, and six (*Obp19*, *Obp63*,*Obp66*, *Obp67*, *Or50*, *Or69*) were reassigned to a different orthologue. Five genes annotated as Obps (*Obp69*, *Obp70*, *Obp71*, *Obp74*, *Obp76*) are D7 salivary proteins, which share the Obp lipocalin fold but are not olfactory. One gene annotated as *Ir7y* is *Ir21a* (AGAP008511). All four of our previous principal anthropophily candidates (*Ir8a*, *Or75*, *Obp22*, Gr57) were confirmed. Each gene was placed on the *An. farauti* FAR1 assembly using its *An. farauti* FAR1 gene ID (AFAF) (*75*). We mapped *An. gambiae* IDs to AFAF IDs through the VectorBase orthology service and took gene boundaries (scaffold, start, end, strand) from the FAR1 annotation. Genes without a VectorBase orthologue were placed by BLASTP of the *An. gambiae* protein against the FAR1 annotated proteome (VectorBase release 68). In total, 206 of the 210 genes received FAR1 coordinates. Candidate variant sites were the variants inside these gene boundaries (no flanking sequence) that our variant-effect annotation (see genetic-load methods) classified as HIGH impact (e.g. stop-gained, stop-lost, splice-site) or MODERATE impact (missense), polarised using the *An. punctulatus* outgroup as the ancestral state as in our genetic-load analysis.

Relate selection scan

For each taxon, we pruned the joint ARG (see above - Relate inference of ARGs) to that taxon's haplotypes (RelateExtract --mode SubTreesForSubpopulation). We then re-estimated branch lengths under the taxon's own coalescence-rate history (*Ne*(t)), estimated genome-wide by pooling all 46 scaffolds (EstimatePopulationSize.sh). These taxon-specific trees were used to produce the selection coefficients and the CLUES2 inference below. A SNP's mutation age is the midpoint of the branch its mutation maps to, (age_begin + age_end)/2, taken from the re-estimated .mut output. Note that the Relate estimate of *Ne* (t) was concordant with the MSMC-IM estimate.

For each of six taxa (*An.* farauti s.s., *farauti 8, hinesorum, irenicus, oreios,* and *torresiensi*s) we ran RelateSelection on the taxon-specific trees. The *An. punctulatus* outgroup was excluded. For each SNP and each of 31 fixed time epochs, this test asks whether the derived allele rose in frequency faster than expected under neutral drift, given the taxon's Ne(t). The one-sided test only detects frequency increases, and so cannot distinguish between purifying selection or selection against the derived allele.

We removed SNPs whose local tree carried fewer than 10 mapped SNPs (num_snps_on_tree < 10, from RelateSelection's .qual output). We called a SNP significant if its minimum p-value across epochs was below a fixed genome-wide threshold of $5 \times 10^{-8}$ (*55, 115*). The scan was restricted to the HIGH- and MODERATE-impact chemosensory sites defined above: 26,169 taxon-by-site tests across the six taxa.

<u>CLUES2 analysis</u>

For candidate sites, we estimated selection coefficients with CLUES2 (*116*). The coefficient fit is conditioned on a supplied population-size history. We ran it in two taxa: *An. farauti s.s.* (279 individuals, 558 haplotypes) and *An. hinesorum* (126 individuals, 252 haplotypes). These are the only taxa with significant sites in the RelateSelection scan. Candidates were HIGH- and MODERATE-impact chemosensory SNPs that were segregating in the focal taxon: 198 sites (farauti_ss 63, hinesorum 135; 16 HIGH, 182 MODERATE). At each site we drew 200 branch-length samples from the taxon-specific trees (SampleBranchLengths.sh --format n, same mutation rate and Ne(t) as above). The extraction window ran from the focal SNP to the next SNP on the scaffold. The samples were converted to CLUES2 coalescence-time input with RelateToCLUES.py. Derived-allele state for each haplotype, and the site's derived-allele frequency in the taxon (--popFreq), were taken from the polarised genotype matrix in the same sample order as the ARG. CLUES2 was given the taxon's Relate Ne(t) (--coal).

We fitted CLUES2 (inference.py) with 100 frequency bins (--df 100) and without computing the posterior allele-frequency trajectory (--noAlleleTraj). Computing the trajectory needed more than 300 GB of memory per site at this depth; the flag does not affect the estimate of s. The analysis window (--tCutoff) was set separately for each site from its mutation age: tCutoff = max(20,000, 1.5 x age_end) generations, rounded up to the nearest 1,000 generations. We scaled each window to each mutation's age (estimated in Relate): median mutational age is ~2.9 kya and the oldest is ~14 kya. Sites that still returned a boundary value (logLR > $10^6$) were treated as unconverged and excluded. Three sites whose coalescence-time input files were malformed were also excluded. Selection was called significant at the same fixed threshold as the RelateSelection scan, $p < 5 \times 10^{-8}$.

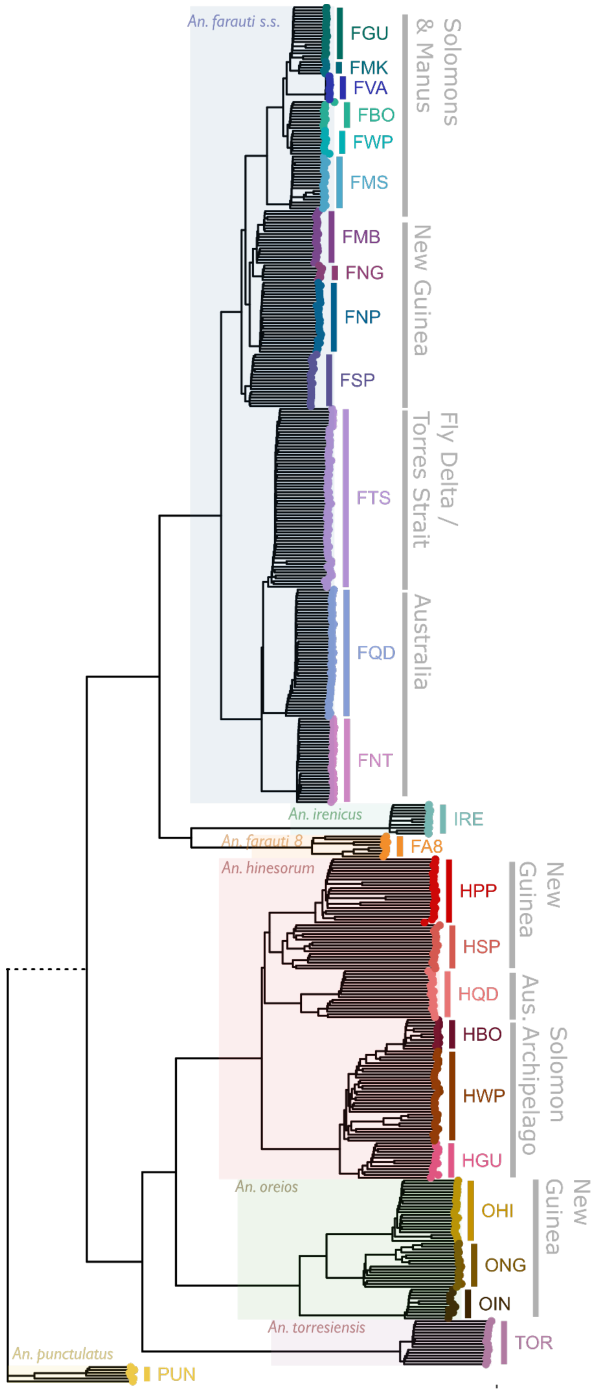


**Fig. S1.**

Neighbour joining tree based on 100,000 randomly chosen SNPs from across the *An. farauti* genome. Shaded boxes indicate taxon (labelled), tip colours and labels indicate population (see Fig. 1B, table S2). Major geographic groups are labelled in grey.

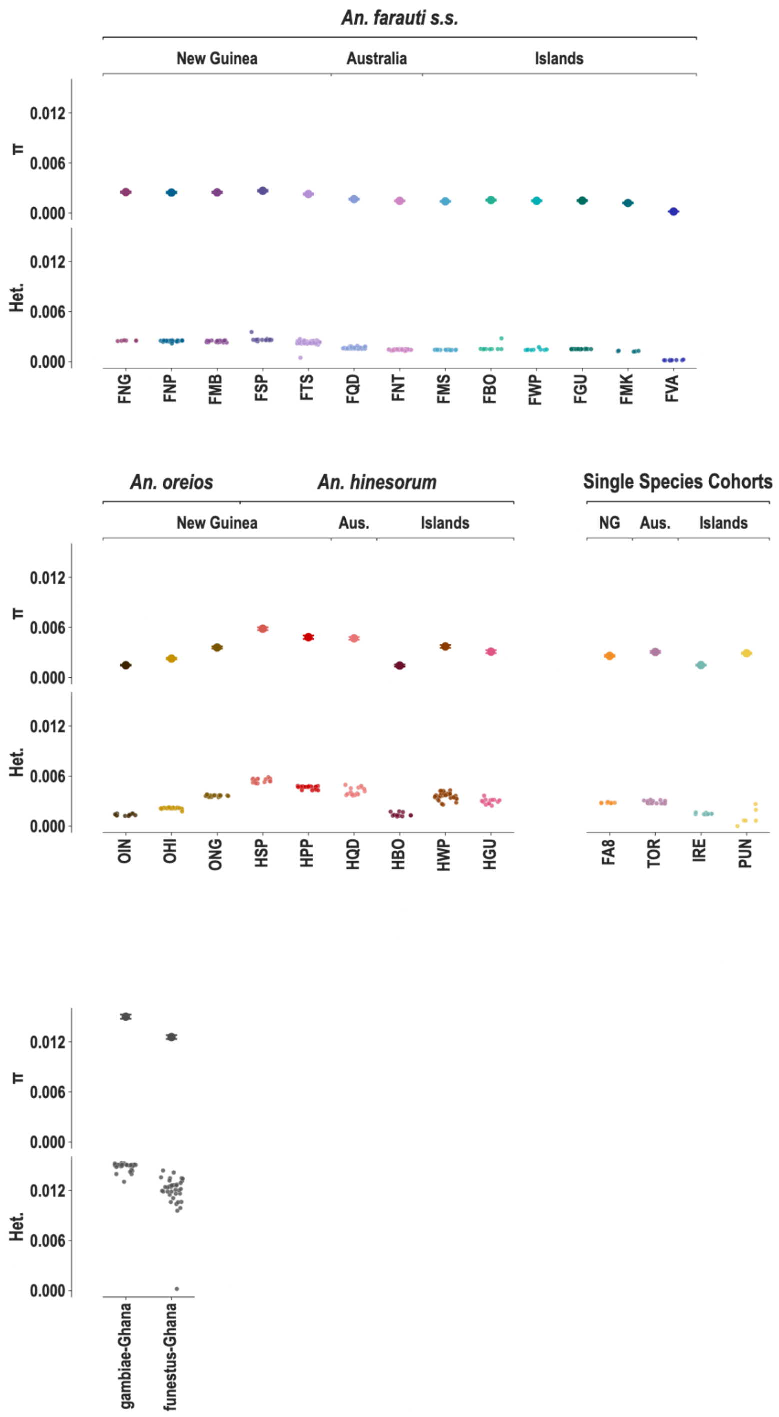


**Fig. S2.**

Genomic diversity of the *An. farauti* complex. Top rows show nucleotide diversity (π, error bars show block-jackknife SE) and per-sample heterozygosity (Het.), grouped by population, across all taxon groups (plot panels). Population labels are shown in full (see table S1 and Fig. 1 for more information). Taxon (labelled across the top), with major geographic regions indicated in subheadings.

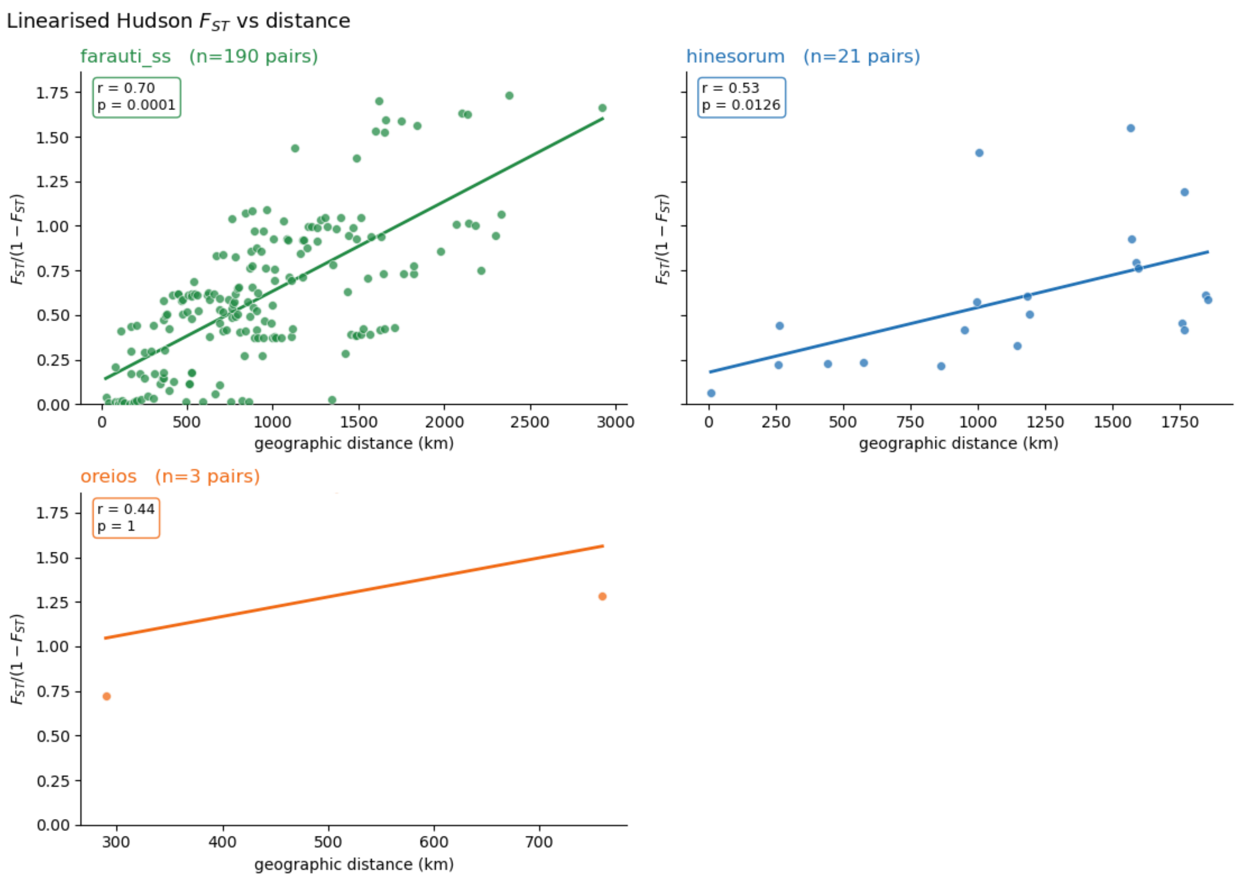


**Fig. S3.**

Strong isolation by distance in *An. farauti ss* and *An. hinesorum*. Plots show linearised Hudson's *Fst* (y axis) plotted against geographic distance in km (x axis), for *An. farauti ss* (left, green) and *An. hinesorum* (right, blue), computed between sampling sites with >5 samples per location as per((*36*)). Regression lines are plotted, with model statistics enclosed on the top left of plots.

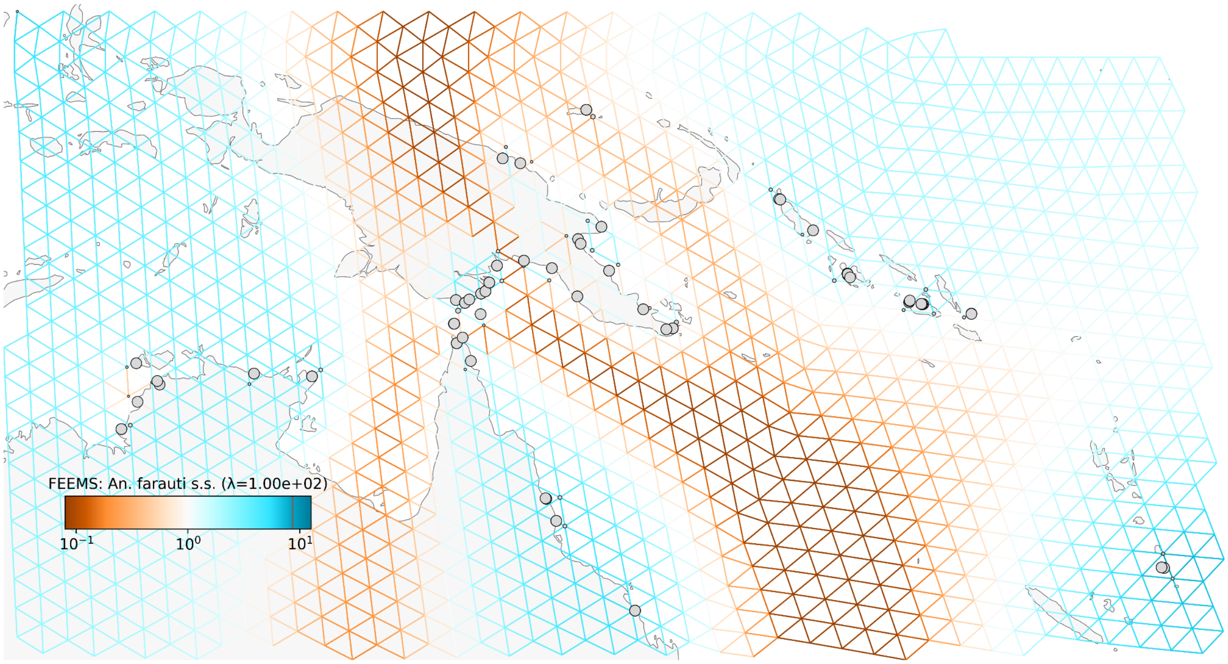


**Fig. S4.**

fEEMS - *An. farauti*. The grid shows inferred effective migration ($\log_{10}(w)$) for *An. farauti* s.s. across Australia, PNG, Solomon Islands and Vanuatu estimated from *fEEMS*. The grid is coloured from dark brown (low) to bright blue (high) between sampling nodes (grey points).

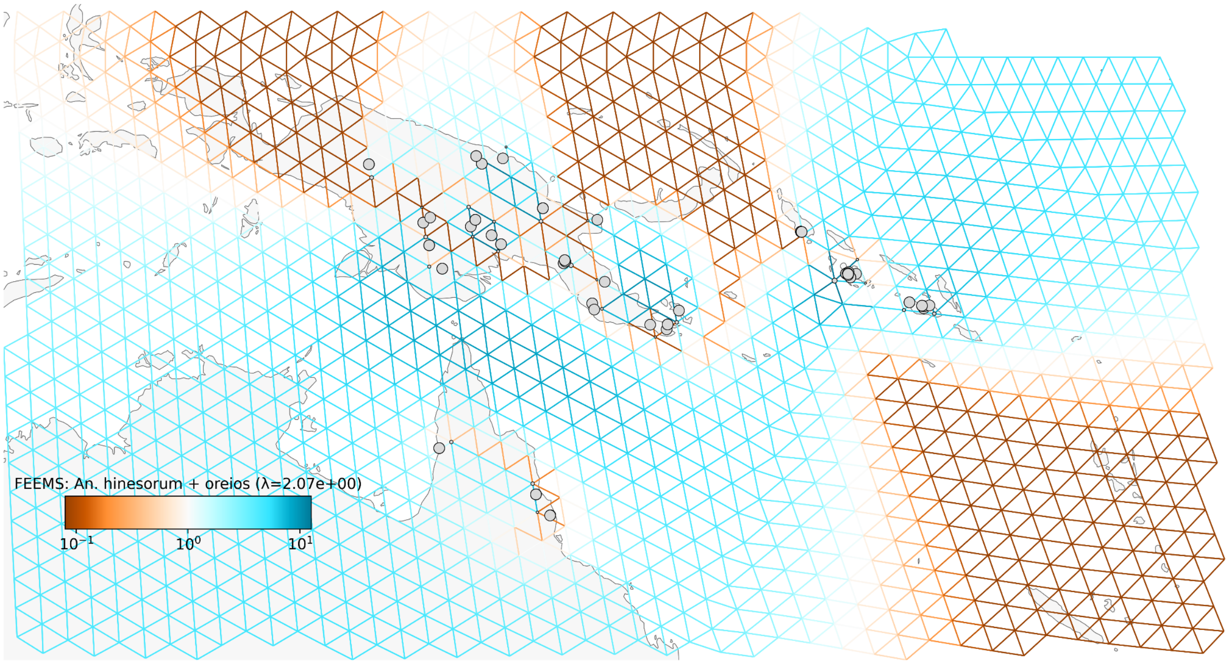


**Fig. S5.**

fEEMS - *An. hinesorum* and *An. oreios.* The grid shows inferred effective migration ($\log_{10}$(w)) for *An. farauti ss* and *An. hinesorum* across Australia, PNG, Solomon Islands and Vanuatu estimated from *fEEMS*. The grid is coloured from dark brown (low) to bright blue (high) between sampling nodes (grey points).

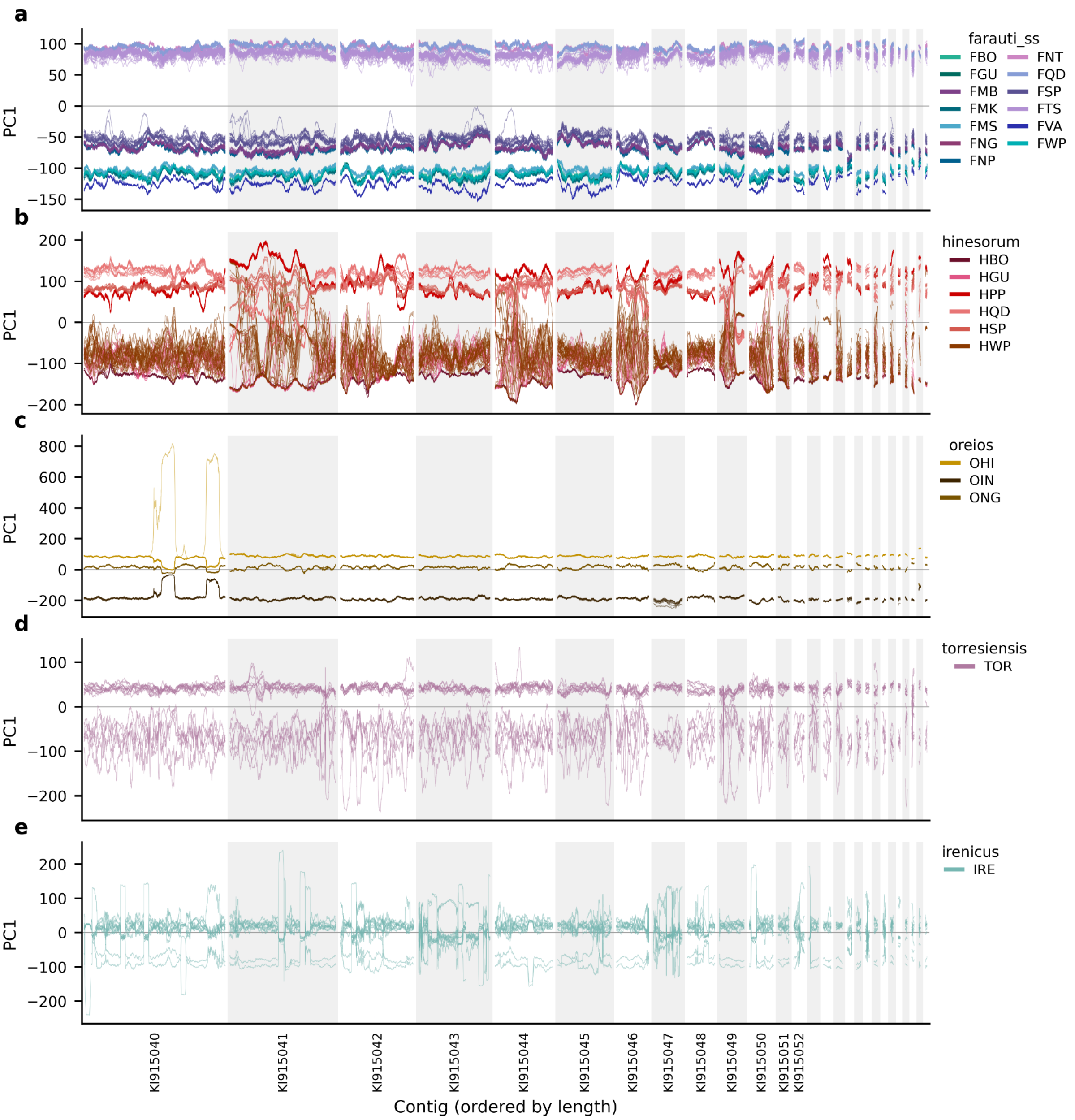


**Fig. S6.**

PC1 (y axis) of principal component analysis (PCAs) per-taxon (plot rows) calculated using accessible SNPs in 50kb windows for all samples across all contigs in the *An. farauti* genome. Alternate shading shows contigs. Individual lines indicate samples, coloured by population (see keys on right hand side). In the presence of an inversion polymorphism, PCA will usually reflect the karyotype (e.g. homozygous negative, heterozygous, homozygous positive), as opposed to the neutral structure of the population((*99*), (*33*)). The plots above show an absence of structuring by inversion karyotype.

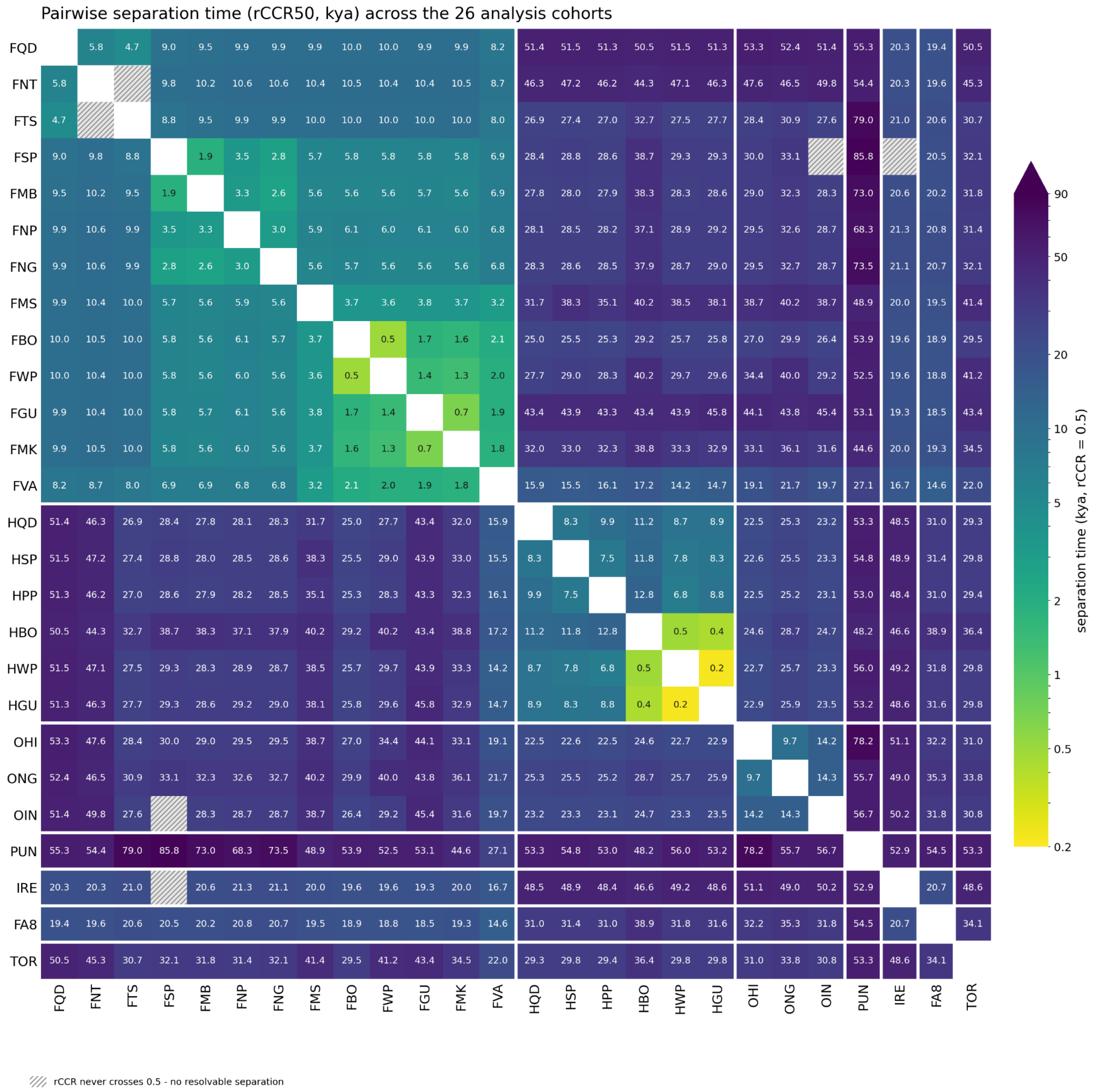


**Fig. S7.**

Estimated pairwise population separation times between analysis populations. Each cell contains a point estimate of population separation time, based on the M(t) statistic generated by MSMC-IM (see Methods). The M(t) indicates when the cumulative migration probability, based on relative cross-coalescence rate, reaches 0.5. Cell heat is scaled by M(t). The diagonal is zeroed. Cells containing hashed lines indicate pairs for which the algorithm did not converge.

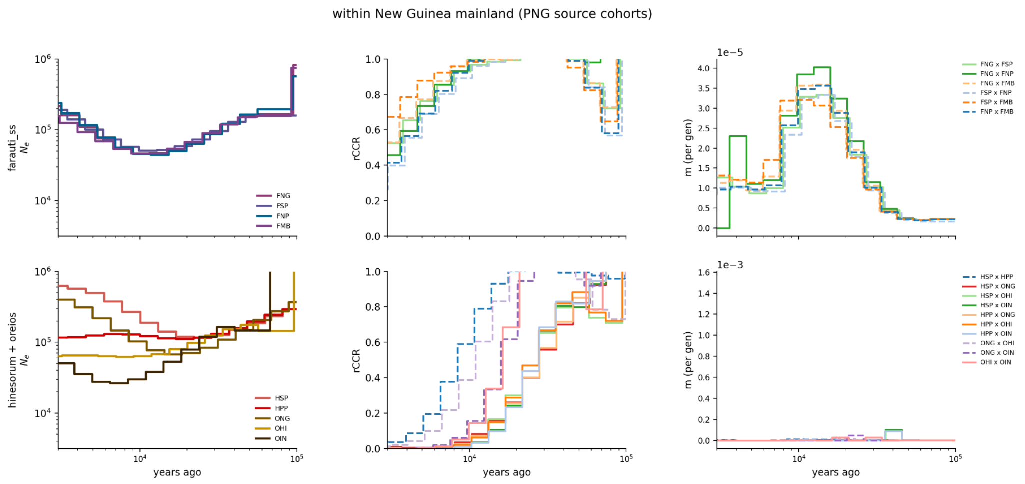


**Fig. S8.**

Within population demographic history and gene flow for the New Guinea mainland populations. Left: MSMC2 estimates of effective population size (*Ne*), middle: relative cross-coalescence rate (rCCR) and right: symmetric migration rate (m, per generation), over the last 4000-100,000 years. Top row: *An. farauti s.s.* populations, bottom: *An. hinesorum* and *An. oreios* populations, for every population combination within each taxon group for this region, coloured/styled as per the legends. Split times discussed in the text are taken as the point at which rCCR crosses 0.5. Note that the last 4,000 years have been truncated from the x axis, as per Methods.

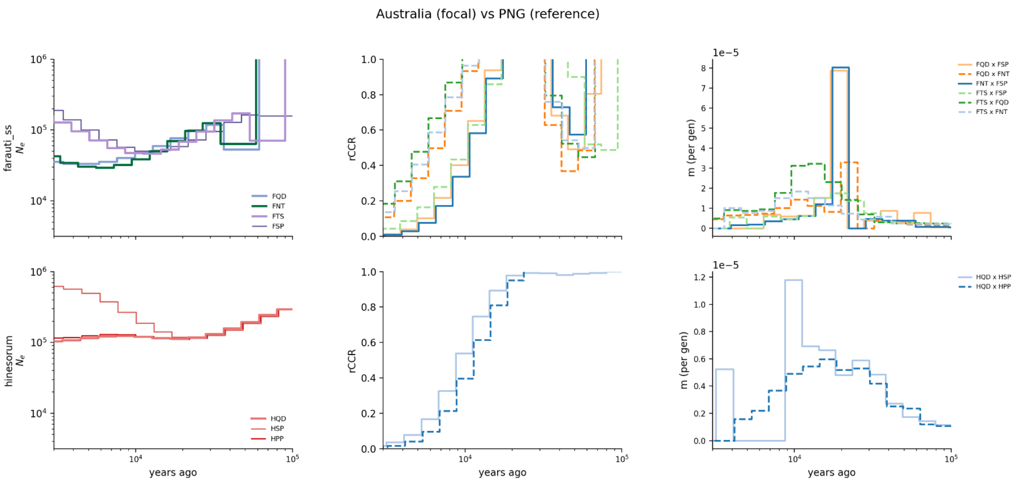


**Fig. S9.**

Within population demographic history and gene flow for the Australian vs New Guinea populations. Left: MSMC2 estimates of effective population size (*Ne*), middle: relative cross-coalescence rate (rCCR) and right: symmetric migration rate (m, per generation), over the last 4000-100,000 years. Top row: *An. farauti s.s.* populations, bottom: *An. hinesorum* populations, for every population combination within each taxon group for this region, coloured/styled as per the legends. Split times discussed in the text are taken as the point at which rCCR crosses 0.5. Note that the last 4,000 years have been truncated from the x axis, as per Methods.

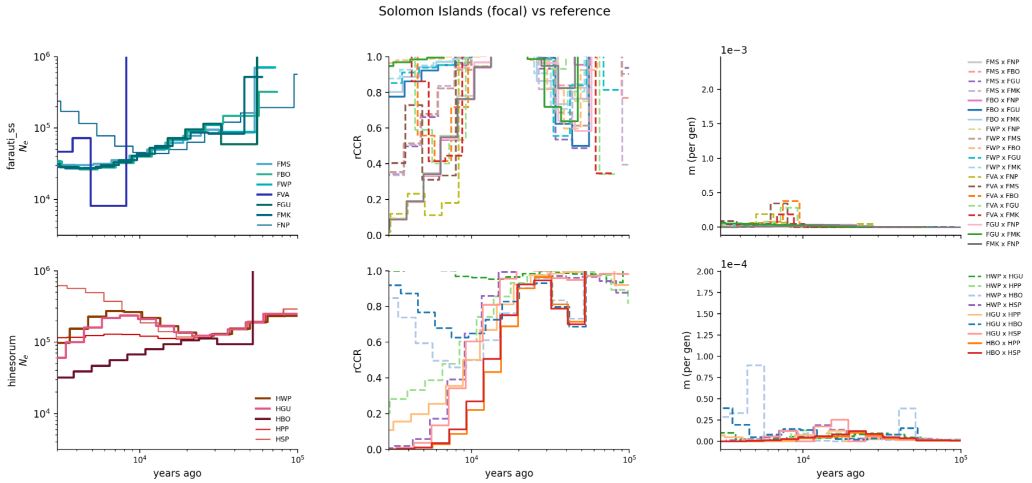


**Fig. S10.**

Within population demographic history and gene flow for the Solomon Islands vs New Guinea populations. Left: MSMC2 estimates of effective population size (*Ne*), middle: relative cross-coalescence rate (rCCR) and right: symmetric migration rate (m, per generation), over the last 4000-100,000 years. Top row: *An. farauti s.s.* populations, bottom: *An. hinesorum* populations, for every population combination within each taxon group for this region, coloured/styled as per the legends. Split times discussed in the text are taken as the point at which rCCR crosses 0.5. For *An. farauti,* the northern Papuan Peninsula population (FNP) was used as a reference population; for *An. hinesorum:* Papuan peninsula (FPP) and southern New Guinea (HSP). Note that the last 4,000 years have been truncated from the x axis, as per Methods.

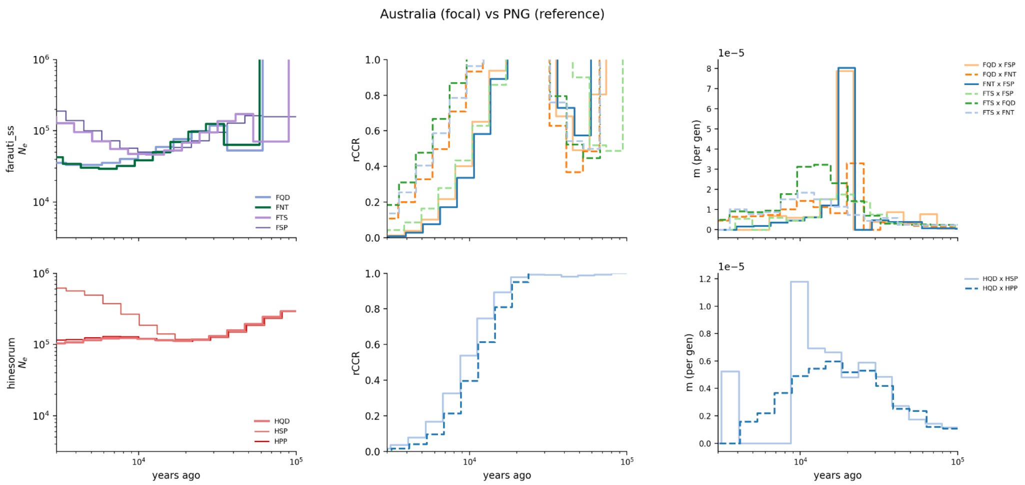


**Fig. S11.**

Within population demographic history and gene flow for the Australian vs New Guinea populations. Left: MSMC2 estimates of effective population size (*Ne*), middle: relative cross-coalescence rate (rCCR) and right: symmetric migration rate (m, per generation), over the last 4000-100,000 years. Top row: *An. farauti s.s.* populations, bottom: *An. hinesorum* populations, for every population combination within each taxon group for this region, coloured/styled as per the legends. Split times discussed in the text are taken as the point at which rCCR crosses 0.5. For *An. farauti,* the northern Papuan Peninsula population (FNP) was used as a reference population; for *An. hinesorum:* Papuan peninsula (FPP) and southern New Guinea (HSP). Note that the last 4,000 years have been truncated from the x axis, as per Methods.

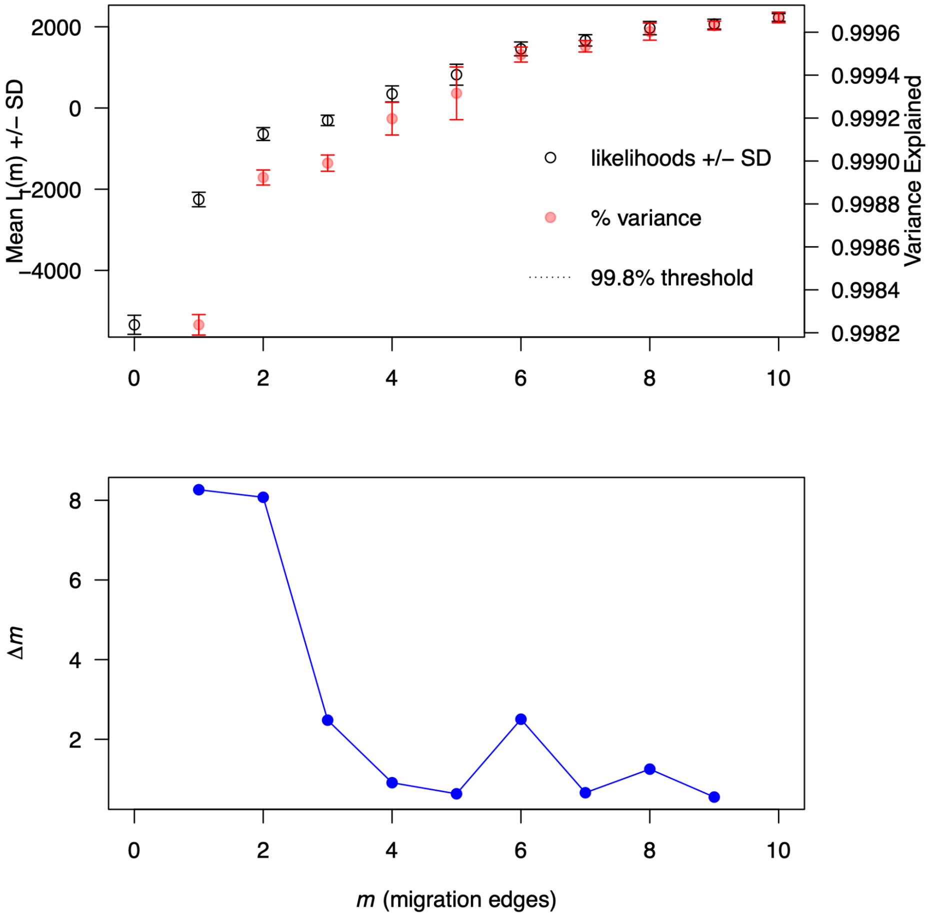


**Fig. S12.**

Changes in the mean likelihood score (± SD) and the average total fraction of the genetic variance explained (± SD) in function of the number of migration edges considered in the *TreeMix* analysis (upper panel). Second-order rate of change in likelihood (Δm) across migration edges (m) values, from 0 to 10 (lower panel). The two first highest rates of likelihood changes are observed for m=1 or m=2, which indicate that migration and admixture are present in the tree. The optimal m-value is observed at m=6.

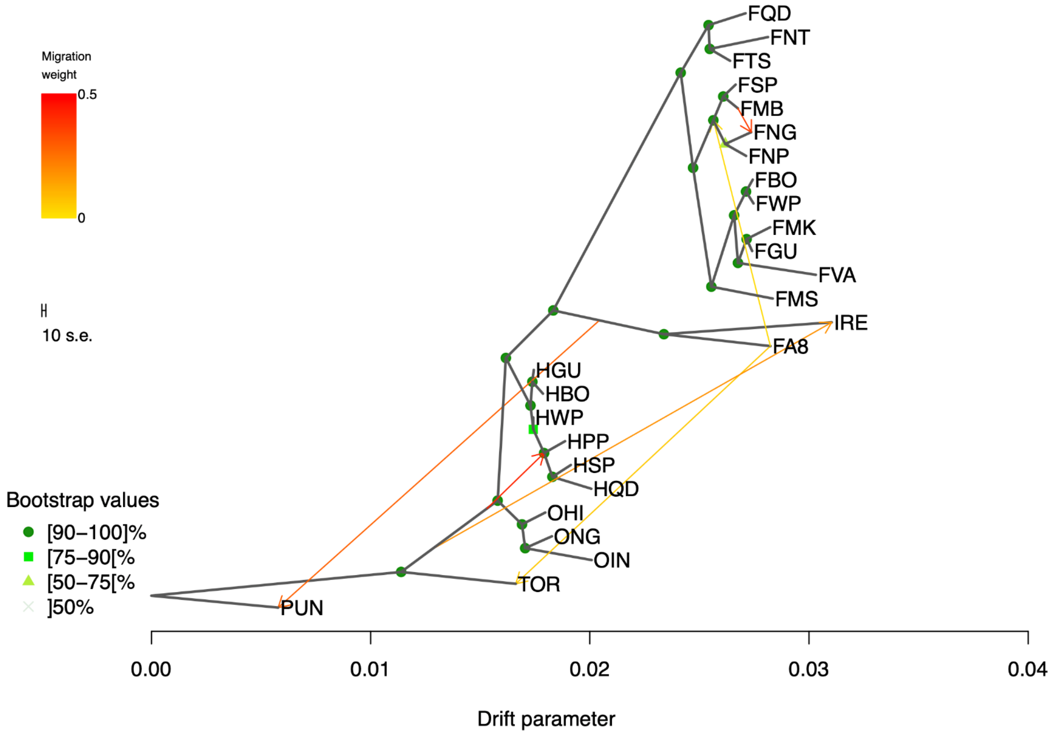


**Fig. S13.**

Most likely population graph topology recovered with *TreeMix* with 6 predicted migration edges. The graph is rooted with An. punctulatus (PUN). Nodes' supports are indicated with the colored circle at the node. The strength of migration / admixture is indicated by the color of the arrow from yellow (light migration) to red (strong migration).

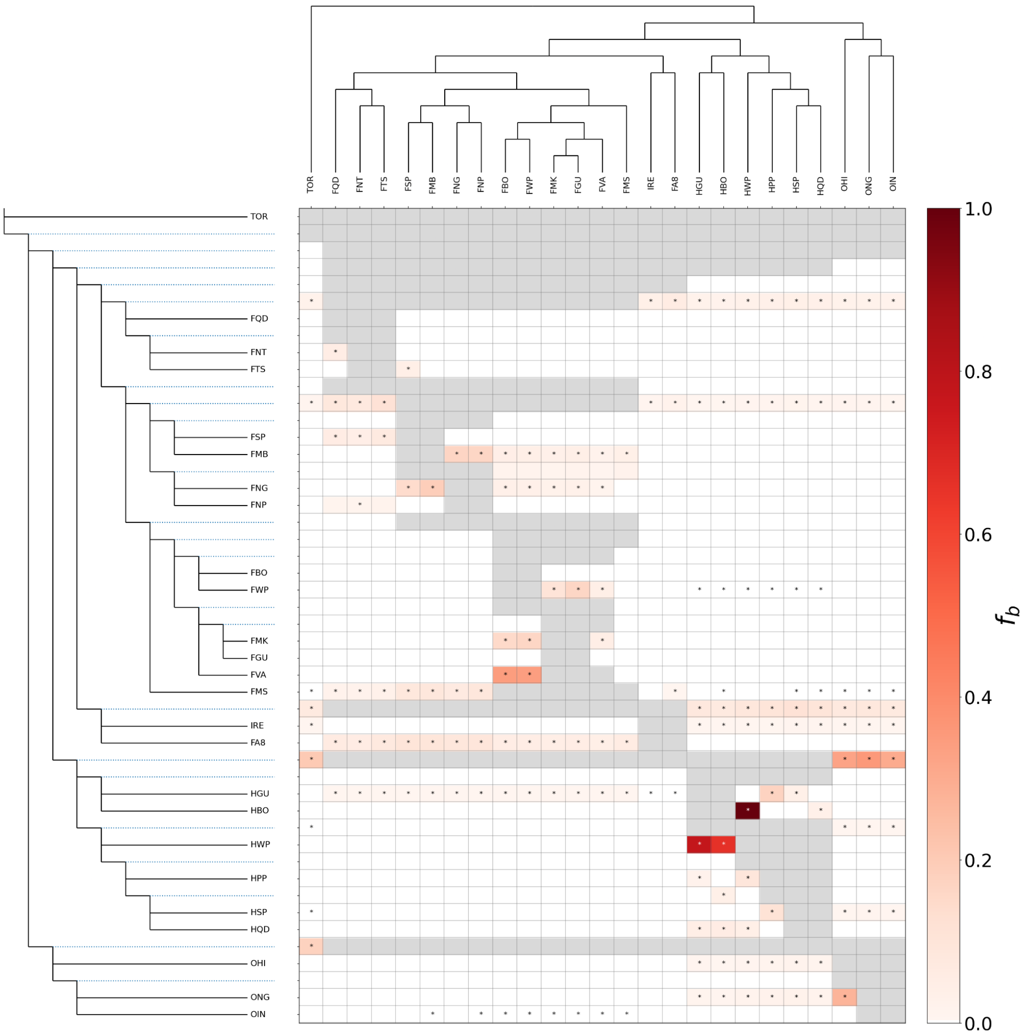


**Fig. S14.**

Branch-specific excess allele sharing across the farauti complex (Dsuite f-branch). The left tree is the population-level species consensus tree (from the best-fitting TreeMix analysis) used to arrange the trios; every branch, terminal and internal, is a row. Dotted blue lines link each internal branch to its row. The top tree shows the same topology for the 25 ingroup populations, one per column. Each cell gives the f-branch statistic $f_b(C)$, the excess allele sharing between the row branch b and the column population C. It is read as a rough proportion of shared ancestry beyond what the tree predicts. Values were computed from all 2,300 population trios. Dtrios was run on 480 individuals across 46 genomic contigs and the per-contig results combined with DtriosCombine, with *An. punctulatus* (n = 6) as the outgroup. f4-ratio values with $p >= 0.01$ were set to zero before summarising (Dsuite fbranch default). Grey cells are branch-population pairs that the tree topology leaves untestable, for example a population paired with its own branch or ancestors. White cells show no significant excess sharing. Asterisks indicate pairs with a significant (*Z*-score >3) amount of excess allele sharing.

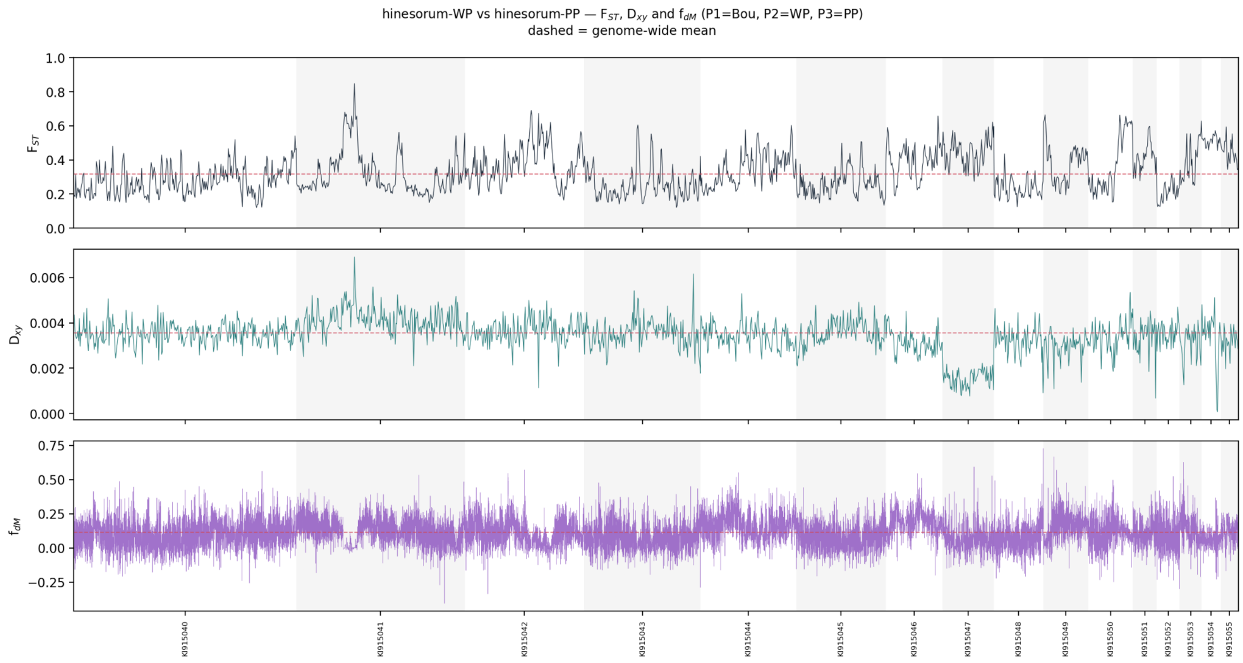


**Fig. S15.**

Genomic scans of differentiation, divergence and introgression between Solomon Islands and PNG *An. hinesorum*. Windowed statistics for hinesorum-WP against hinesorum-PP along the 16 largest scaffolds ordered by decreasing length and shaded alternately. Top: Hudson's $F_{ST}$ in non-overlapping 100 kb windows. Middle: absolute divergence ($D_{xy}$) in the same windows. Both were computed from biallelic segregating sites passing site filters, and windows with fewer than 100 such sites were dropped (1,575 windows retained). Bottom: $f_{dM}$ from Dsuite Dinvestigate for the trio P1 = hinesorum-Bou (n = 11), P2 = hinesorum-WP, P3 = hinesorum-PP, with An. punctulatus (n = 6) as the outgroup, in sliding windows of 50 informative SNPs with a 25-SNP step (41,887 windows, median span 6.8 kb). Positive $f_{dM}$ indicates excess allele sharing between hinesorum-PP and hinesorum-WP; negative values indicate excess sharing between hinesorum-PP and hinesorum-Bou. Dashed red lines mark the genome-wide mean of each statistic: $F_{ST}$ = 0.32 and $D_{xy}$ = 0.0036 (both weighted by sites per window), and $f_{dM}$ = 0.11 (unweighted). $F_{ST}$ ranges from 0.12 to 0.85 across windows. The highest $F_{ST}$ peak, on KI915041, coincides with the highest $D_{xy}$ in the scan and with $f_{dM}$ near zero. Most of KI915047 shows $D_{xy}$ at roughly half the genome-wide mean, with $F_{ST}$ above the mean.

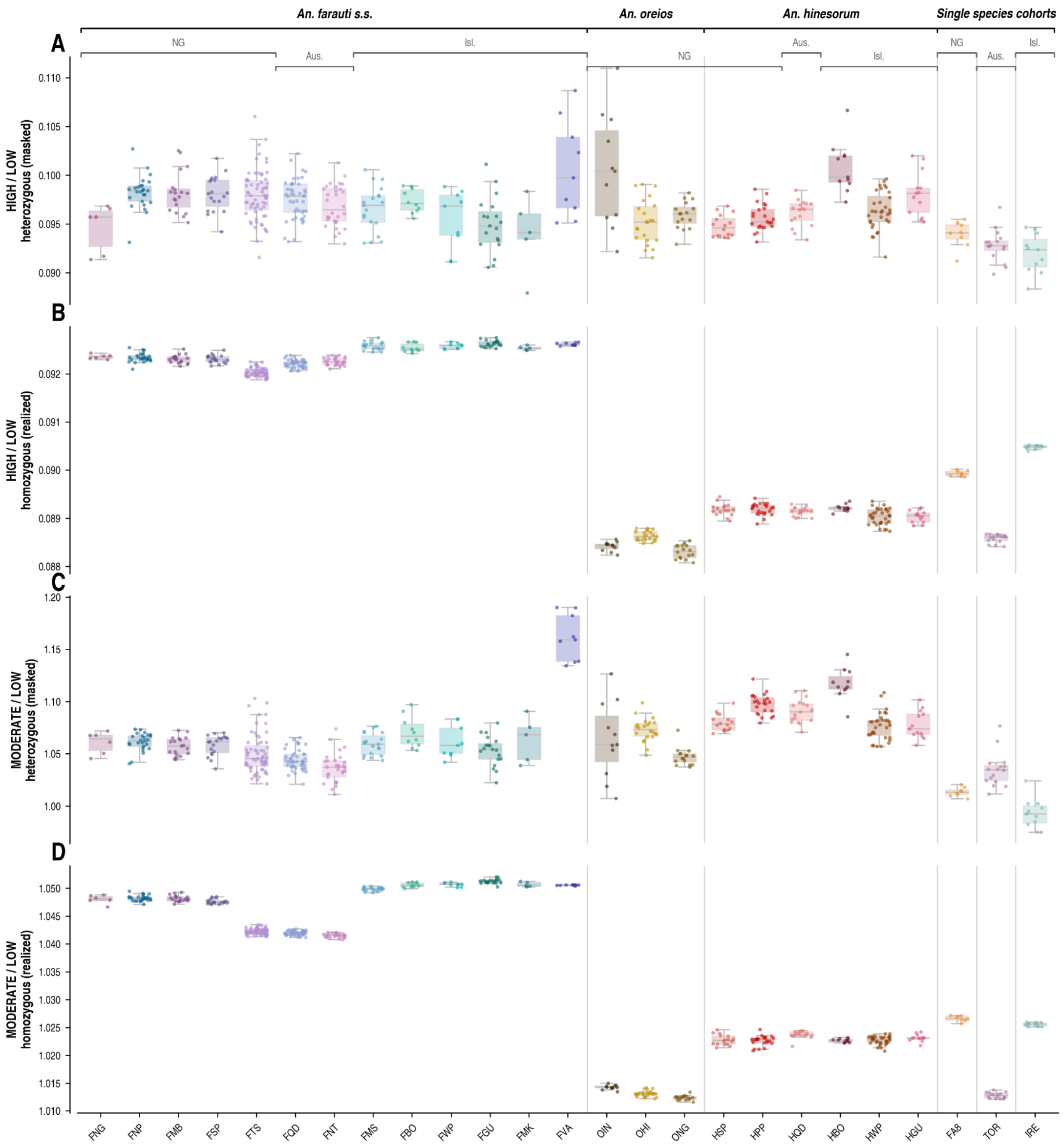


**Fig. S16.** Fully labelled per-sample mutational load for every population (HIGH/LOW and MODERATE/LOW derived-allele ratios, masked and realised load – the labelled version of Fig. 3A–B.

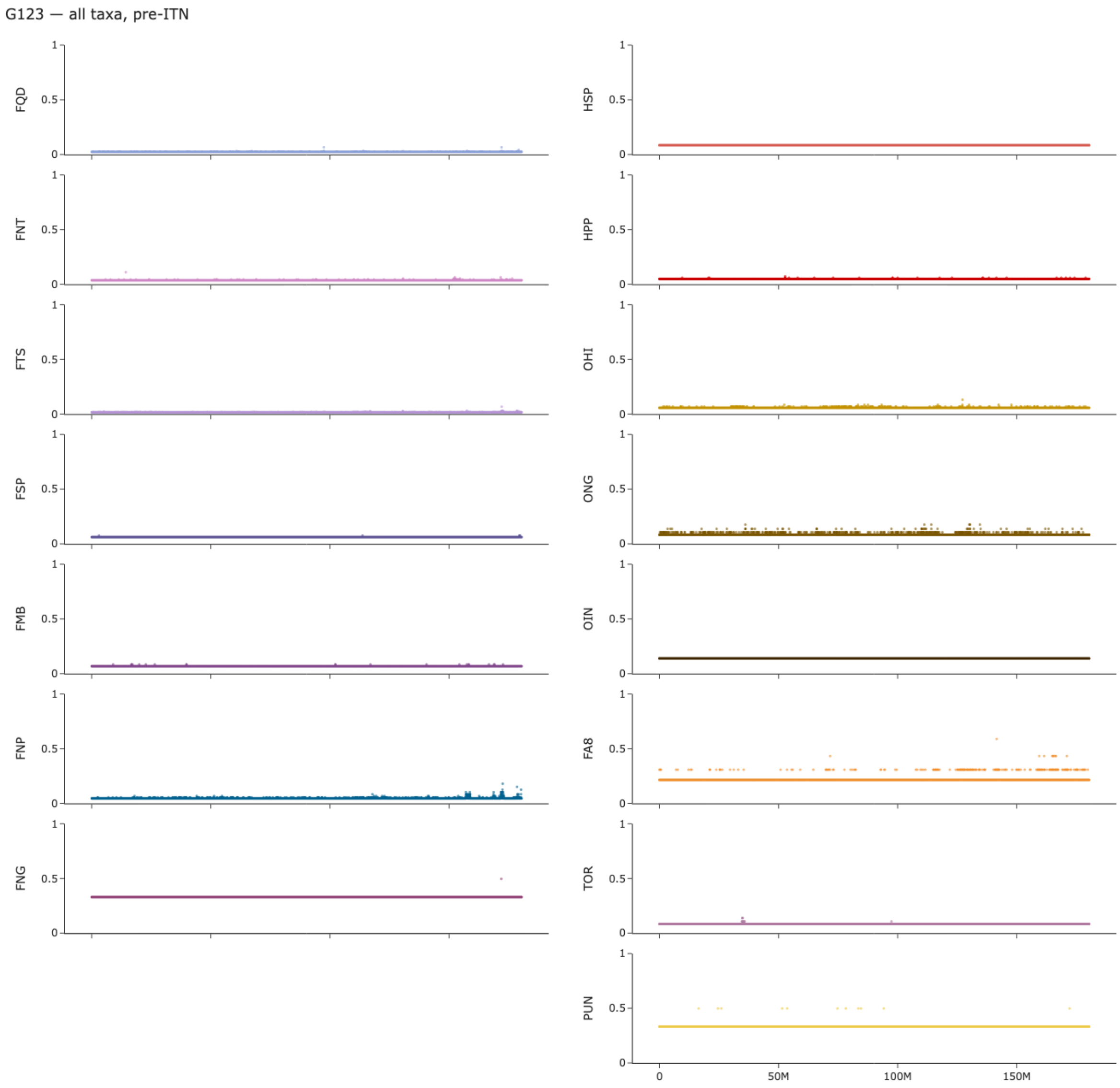


**Fig. S17.**

Genomewide selection scans (GWSS) of *An. farauti* complex populations pre-ITN. Y axis denotes the G123 selection statistic, with the x axis showing the cumulative genomic position across all populations. Plots are panelled by population (Fig. 1, table S2). All populations were sampled between 1990 and 2008: prior to the ITN rollouts of the 2000s, but post the DDT IRS campaigns across the region in the mid-20th century. These populations, with the exception of the island *An. punctulatus,* originate from the ADF surveys in Australia and New Guinea((*24*), (*23*)).

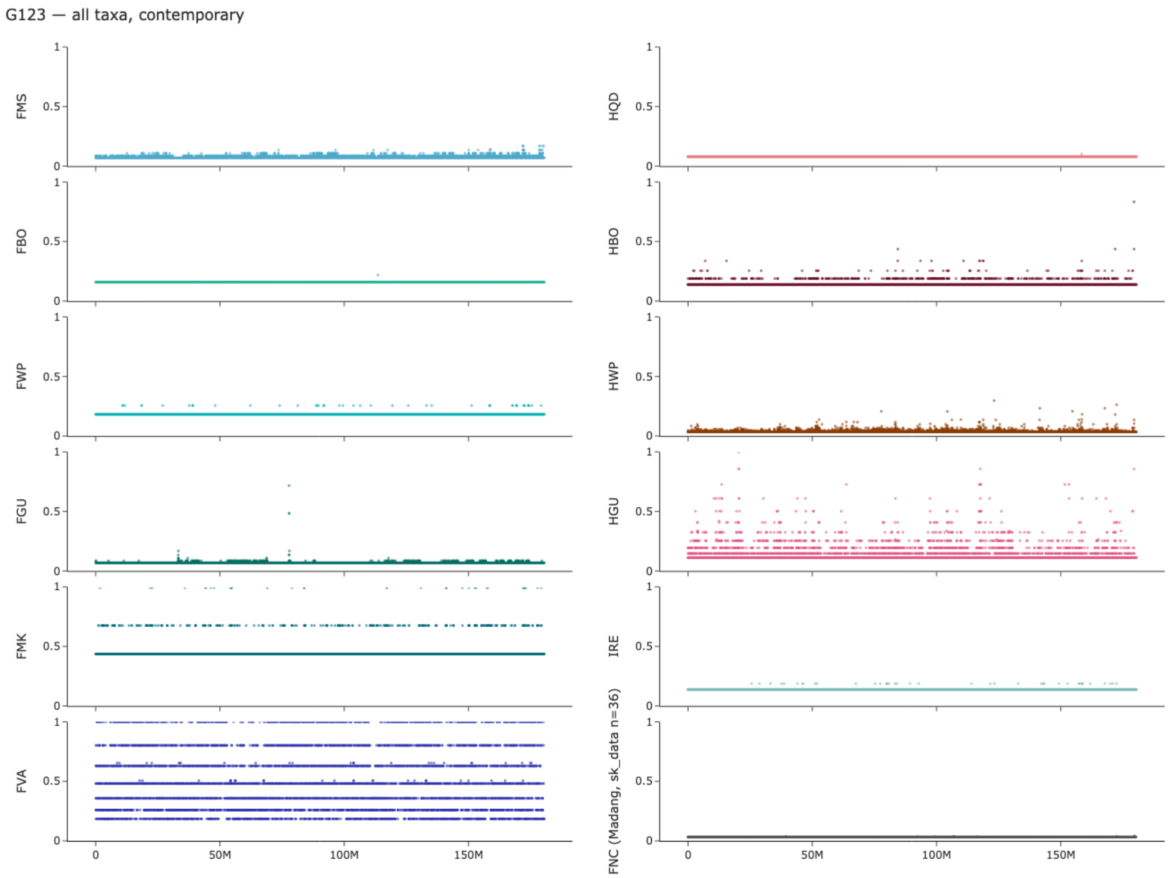


**Fig. S18.**

Genomewide selection scans (GWSS) of *An. farauti* complex populations post-ITN. The Y axis denotes the G123 selection statistic, with the x axis showing the cumulative genomic position across all populations. Plots are panelled by population (Fig. 1, table S2). All populations were sampled post the ITN rollouts of the 2000s. Increased genomewide G123 is likely due to increased bottlenecking and autozygosity of island populations These populations, largely originate from island populations all sampled post the ITN rollouts in the 2000 (see Methods). The only visible sweep is at the *Gste2* locus in *An. farauti s.s.* from Guadalcanal (FGU). Note we added an additional population of samples from northern PNG (FNC, from Madang, bottom right).

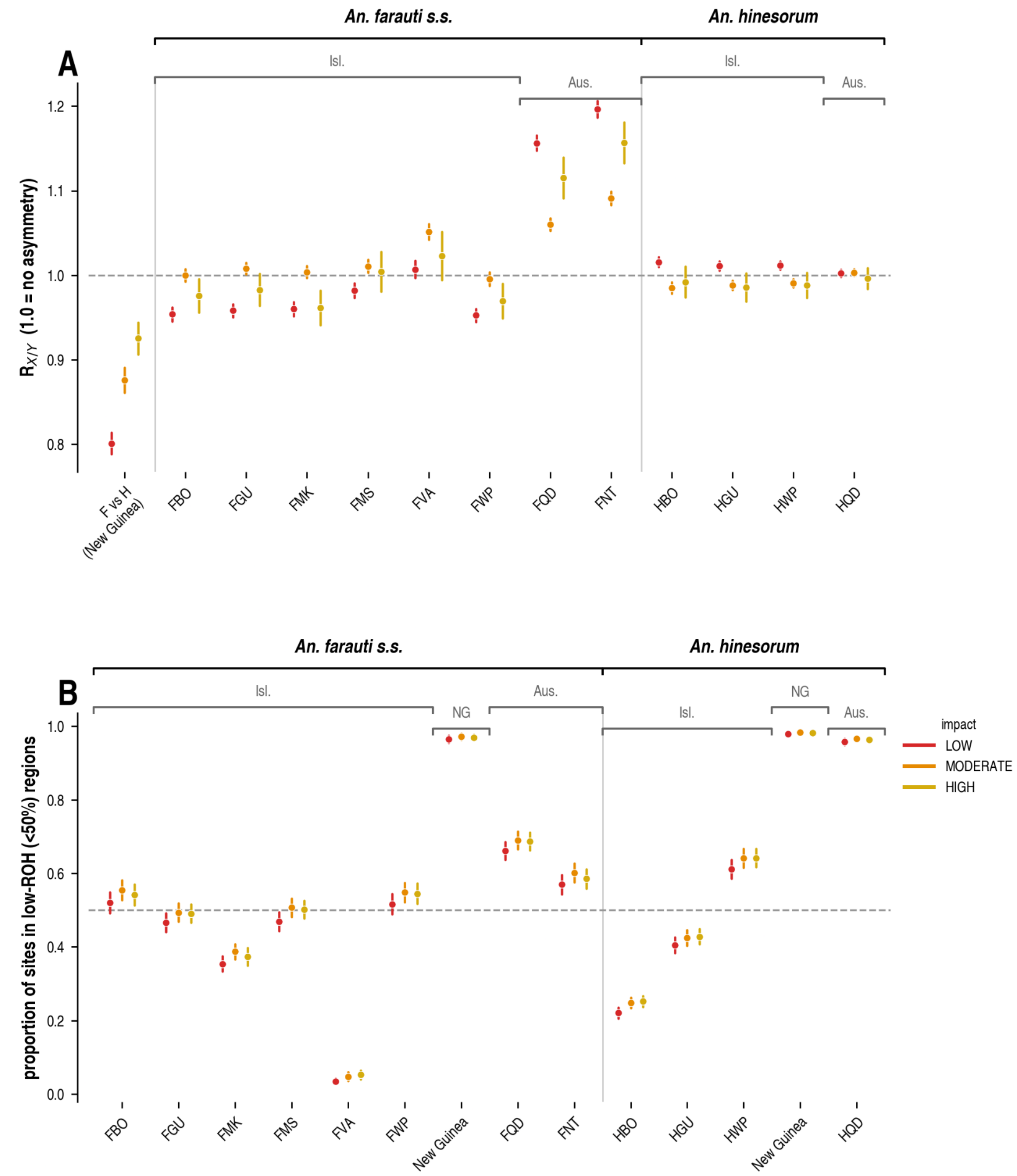


**Fig. S19.** $R_{xy}$ relative deleterious load (A) and proportion of segregating sites in low ROH regions (B) by population – the labelled version of Fig. 3C-D.
